# Impact of 100 LRRK2 variants linked to Parkinson’s Disease on kinase activity and microtubule binding

**DOI:** 10.1101/2022.04.01.486724

**Authors:** Alexia F Kalogeropulou, Elena Purlyte, Francesca Tonelli, Sven M Lange, Melanie Wightman, Alan R Prescott, Shalini Padmanabhan, Esther Sammler, Dario R Alessi

## Abstract

Mutations enhancing the kinase activity of LRRK2 cause Parkinson’s disease (PD) and therapies that reduce LRRK2 kinase activity are being tested in clinical trials. Numerous rare variants of unknown clinical significance have been reported, but how the vast majority impact on LRRK2 function is unknown. Here, we investigate 100 LRRK2 variants linked to PD, including previously described pathogenic mutations. We identify 23 LRRK2 variants that robustly stimulate kinase activity, including variants within the N-terminal non-catalytic regions [ARM (E334K, A419V), ANK(R767H), LRR (R1067Q, R1325Q)], as well as variants predicted to destabilise the ROC:COR_B_ interface [ROC (A1442P, V1447M), COR_A_ (R1628P) COR_B_ (S1761R, L1795F)] and COR:COR dimer interface [COR_B_ (R1728H/L)]. Most activating variants decrease LRRK2 biomarker site phosphorylation (pSer935/pSer955/pSer973), consistent with the notion that the active kinase conformation blocks their phosphorylation. We conclude that the impact of variants on kinase activity is best evaluated by deploying a cellular assay of LRRK2-dependent Rab10 substrate phosphorylation, compared to a biochemical kinase assay, as only a minority of activating variants [COR_B_ (Y1699C, R1728H/L, S1761R) and kinase (G2019S, I2020T, T2031S)], enhance *in vitro* kinase activity of immunoprecipitated LRRK2. Twelve variants including several that activate LRRK2 and have been linked to PD, suppressed microtubule association in the presence of a Type I kinase inhibitor [ARM(M712V), LRR(R1320S), ROC (A1442P, K1468E, S1508R), COR_A_(A1589S), COR_B_ (Y1699C, R1728H/L) and WD40(R2143M, S2350I, G2385R)]. Our findings will stimulate work to better understand the mechanisms by which variants impact biology and provide rationale for variant carrier inclusion or exclusion in ongoing and future LRRK2 inhibitor clinical trials.

## Introduction

1-4% of all cases of PD are caused by genetic changes in Leucine-Rich Repeat Kinase-2 (LRRK2) [1–3]. Additionally, LRRK2 has been linked to modify risk for Crohn’s disease (CD) [4]. LRRK2 is a large multidomain enzyme that forms multimeric species [5, 6]. It consists of an N-terminus armadillo (ARM), ankyrin (ANK) and leucine-rich repeats (LRR), followed by a C-terminal Roco type GTPase, protein kinase and WD40 domain [7] (Figure 1A). The Roco GTPase domain consists of three subdomains, namely a ROC GTPase followed by two scaffolding domains termed COR_A_ and COR_B_. High resolution Cryo-EM structures of full length [8], as well as the catalytic C-terminal moiety of LRRK2, have been solved in which the catalytic kinase domain is in an inactive open conformation, [9] and more recently in a closed conformation [10]. These structures have provided major insights into the overall structure and function of LRRK2.

**Figure 1.**
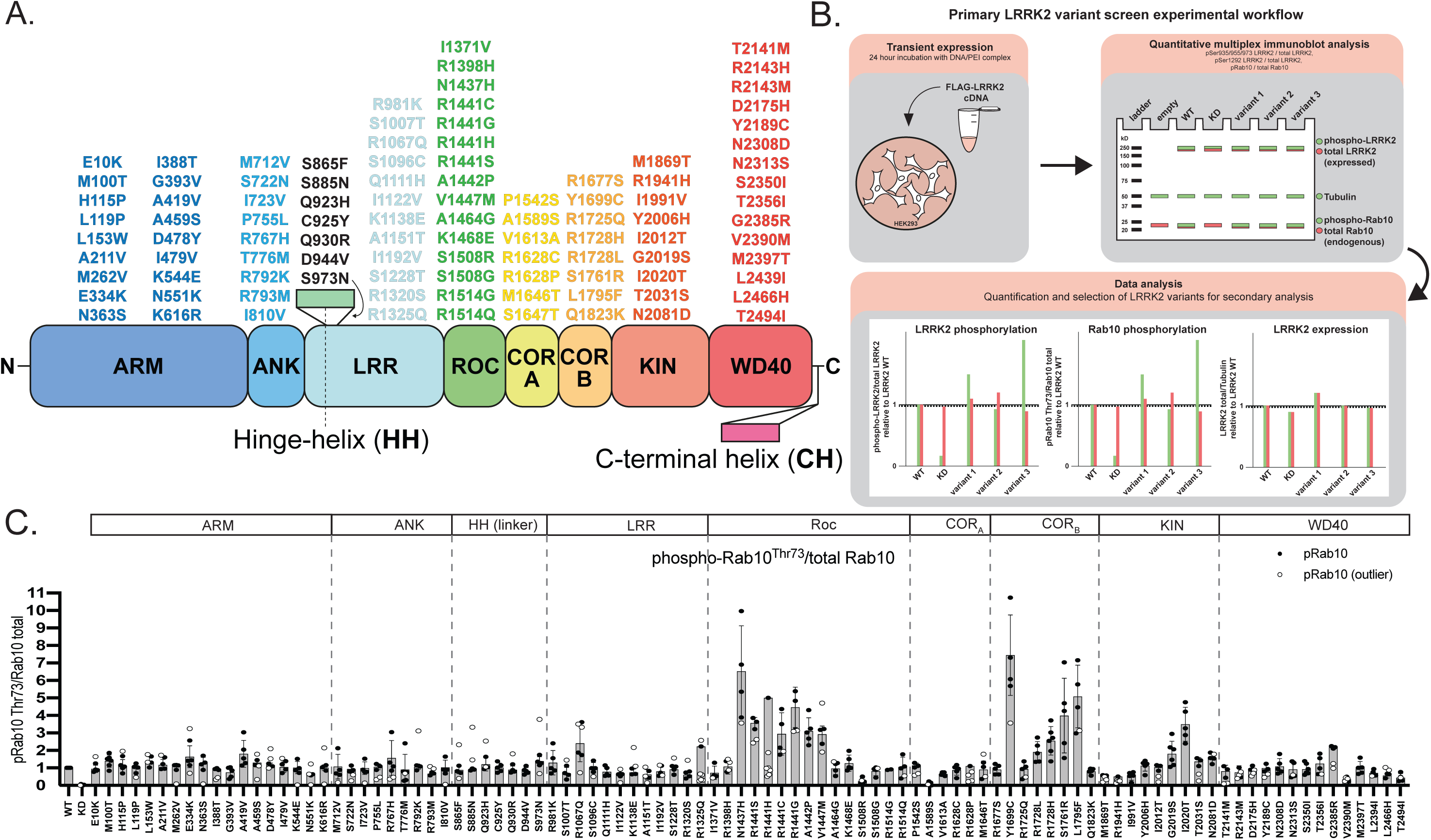
Domain location of 100 LRRK2 variants and experimental workflow to assess LRRK2 variant activity by quantitative immunoblotting. (A) LRRK2 domain structure highlighting 100 PD and CD associated variants within the Armadillo (ARM), Ankyrin (ANK), Leucine Rich Repeats (LRR), Ras of complex proteins (ROC), C-terminal of Roc A and B (COR_A_, COR_B_), Kinase (KIN), and WD40 domains. The LRRK2 variants located in the linker region between the hinge-helix (HH) and LRR domain are listed in black. (B) Workflow schematic outlining the characterisation of the selected LRRK2 variants in a HEK293 overexpression system, followed by quantitative immunoblotting and quantitation of LRRK2 activity relative to wildtype LRRK2. (C) FLAG-tagged LRRK2 wildtype, kinase dead (KD = D2017A), and the indicated variants were transiently expressed in HEK293 cells. 24 hours post-transfection, cells were lysed and analysed by quantitative immunoblotting (as in Supplementary Figure 2A). Quantified immunoblotting data are presented as ratios of pRab10 Thr73/total Rab10, normalized to the average of LRRK2 wildtype values for each replicate (mean ± SD). Combined immunoblotting data from up to 6 independent biological replicates are shown. Dashed lines segment the graphs into corresponding regions of LRRK2 as listed in the domain schematic.

LRRK2 is activated following its recruitment to cellular membranes via interactions with Rab29 and other Rab GTPases, likely via its N-terminal ARM domain [11–15]. LRRK2 phosphorylates a subgroup of Rab GTPases at membranes, including Rab8A and Rab10, at a conserved Ser/Thr residue located within the effector binding Switch-II domain [16–18]. This phosphorylation does not impact intrinsic Rab GTPase activity but promotes binding to a new set of effectors, including RILPL1/2 and JIP3/JIP4 [17,19–22]. Interaction of LRRK2-phosphorylated Rab8A and Rab10 with RILPL1 interferes with ciliogenesis in brain cholinergic neurons in the striatum, decreasing their ability to sense Sonic hedgehog in a neuro-protective circuit that supports dopaminergic neurons [19].

LRRK2-phosphorylated Rab proteins are dephosphorylated by the PPM1H phosphatase [23]. At least in overexpression studies, certain pathogenic mutations, as well as treatment with selective Type I kinase inhibitors that promote the LRRK2 kinase domain to adopt an active conformation, induce helical oligomerization of LRRK2 on microtubule filaments [24–27]. This has been proposed to disrupt vesicle trafficking by causing a “roadblock” for microtubule-based motors [6, 9]. The closed, active conformation of LRRK2 also leads to the dephosphorylation of a cluster of phosphorylation sites (Ser910, Ser935, Ser955 and Ser973) located between the Hinge helix and LRR domain through an unknown mechanism [28–30]. Certain pathogenic mutations such as G2019S (located within the kinase domain) promote autophosphorylation of LRRK2 at Ser1292 [31].

Seven missense mutations located within the ROC (N1437H, R1441G/C/H), COR_B_ (Y1699C), and kinase (G2019S, I2020T) domains have been well-characterized and ascertained to stimulate LRRK2 kinase activity and cause PD [3, 32]. The G2019S mutation that substitutes a glycine for serine within the magnesium-binding DYG motif is by far the most frequent PD-associated LRRK2 mutation [33, 34]. In addition, a variant located within the WD40 domain (G2385R), is common in Chinese Han and Taiwanese populations and moderately increases PD risk; biochemical analysis suggests that G2385R blocks WD40 dimerization and moderately enhances LRRK2 kinase activity [35–37]. Over 1000 rare variants of LRRK2 have been reported [38–40], and a recent study employed a computational tool that reportedly predicts a Parkinson’s pathogenic “REVEL score” for each variant, with a score > 0.600 predicted to be pathogenic [41–43].

Here we describe a robust workflow to experimentally evaluate LRRK2 variant impact on LRRK2 function. Specifically, we utilize LRRK2-dependent Rab10 phosphorylation at Thr73 as a readout for the LRRK2 kinase pathway activity (pRab10^Thr73^), using selective phospho-specific antibodies [44]. From amongst 100 LRRK2 variants, we identified 23 that robustly enhance LRRK2 kinase activity, defined as >1.5-fold above LRRK2 wildtype. These include novel variants within the N-terminal, non-catalytic ARM, ANK and LRR regions, as well as within the ROC and COR_B_ domains, that are predicted to destabilise the interface between the ROC and COR_B_ domains or impact the COR:COR dimer interface. Amongst the 100 variants tested, we also report a subset of 12 variants that suppress the ability of LRRK2 to bind microtubules in the presence of a Type I LRRK2 kinase inhibitor. Overall, our work will assist in the interpretation of the many reported LRRK2 variants of unknown clinical significance identified in individuals and families with PD [41], by informing on variant impact on LRRK2 function and by providing a framework for the thorough functional characterization and cataloguing of other LRRK2 variants of unknown significance. In fact, the functional stratification of LRRK2 variants is particularly important in view of targeted treatments such as LRRK2 kinase inhibitors entering clinical trials.

## Results

### Selection of LRRK2 variants

100 LRRK2 variants were selected from previous genetic analysis of PD patients (STable 1). This list includes the 7 “definitely pathogenic” mutations as listed in the MDSgene database (https://www.mdsgene.org) [ROC (N1437H, R1441 hotspot mutations), COR_B_ (Y1699C), kinase (G2019S, I2020T)] as well as 2 previously characterized variants, including the kinase (T2031S) and WD40 (G2385R) variants that activate LRRK2 (Fig 1A). A variant linked to increased CD disease risk and LRRK2 activation [kinase (N2081D)], as well as variants reported to protect from PD and CD [ARM (N551K) and ROC (R1398H)], were also included [4]. Other than the well characterized variants mentioned above, the remainder have only been reported in a single or small number of cases and studies and often without clear evidence of pathogenicity in line with current guidelines [45]. Literature citations and REVEL scores [41] (http://database.liulab.science/dbNSFP) for each of the selected variants, as well as evolutionary conservation scores for each variant amino acid determined using the Consurf database (https://consurf.tau.ac.il/) [46], are tabulated in STable 1. The selected variants are located within the following domains: ARM (18), ANK (9), LRR (12), ROC (15), COR_A_ (7), COR_B_ (8), kinase (9) and WD40 (15) domains, as well as between the boundaries of the ANK and LRR (5), and LRR and ROC domains (2) (Fig 1A).

### Impact of variants on LRRK2 activity in a cellular assay

To assess the impact of each variant, we utilised a HEK293 cell overexpression system (summarized in Figure 1B) and assessed LRRK2-mediated pRab10^Thr73^, LRRK2 autophosphorylation at Ser1292, as well as LRRK2 biomarker site phosphorylation (Ser935, Ser955 and Ser973). HEK293 cells lend themselves for the interrogation of LRRK2-dependent pRab10^Thr73^ phosphorylation as they have low levels of endogenous LRRK2 but high endogenous Rab10 expression with resulting complete lack of phosphorylation at the LRRK2-dependent Rab10^Thr73^ phospho-site. The wildtype and all LRRK2 variant constructs used in this study contain the common S1647T variant [47]. The S1647T variant is observed in ∼30% of alleles listed on the gnomAD database (280 000 human alleles) [48] and the Parkinson’s disease variant Browser (103 000 human alleles) [49], with no difference between cases and controls. Our data reveal that the T1647 compared to S1647 variant does not impact LRRK2 activity in wildtype, R1441G and G2019S backgrounds. (SFig 1).

In a primary screen, the selected variants were analysed in parallel and normalized to the effect of the LRRK2 wildtype protein, and data were merged from up to 6 independent screens (Fig 1C, 2, SFig 2, SFig 3). For each immunoblot analysis, we also included LRRK2 wildtype for normalization and a kinase inactive LRRK2[D2017A] variant as a negative control for kinase activity. In an additional experiment, we performed a screen with 98 variants treated ± MLi-2 LRRK2 inhibitor [50] and found that this compound suppressed Rab10 phosphorylation in all cases, thereby demonstrating that activity measured is indeed mediated by LRRK2 (SFig 4). LRRK2 variant impact on LRRK2 kinase activity was defined as ‘activating’ if pRab10 ^Thr73^ levels were >1.5-fold and ‘reduced’ if pRab10 ^Thr73^ levels were <0.5-fold relative to the LRRK2 wildtype protein. Our analysis highlighted 23 variants [ARM (E334K, A419V), ANK (R767H), LRR (R981K, R1067Q), boundary between LRR and ROC (R1325Q), ROC (**N1437H**, **R1441G/C/H/S**, A1442P, V1447M), COR_B_ (**Y1699C**, **R1728H**/L, S1761R, L1795F), kinase (**G2019S**, **I2020T**, **T2031S**, **N2081D**) and WD40 (**G2385R**)], that enhance LRRK2-mediated Rab10 phosphorylation (Fig 1C, SFig 2). Twelve of these variants (shown in bold) had previously been reported to stimulate LRRK2 kinase activity. Using the same overexpression system, we then reanalysed 23 of these activating variants from the primary screen in a secondary quantitative immunoblot analysis, in which we confirmed 22 of the 23 variants to robustly enhance LRRK2-mediated Rab10^Thr73^ phosphorylation >1.5-fold above wildtype (Fig 3). Only the LRRK2 R981K variant fell below the 1.5-fold cut-off and could not be confirmed to be activating in the secondary screen (Fig 3). The activity of the R1628P variant which resides in the COR_A_ domain was analysed separately from the group of other activating variants and was found to stimulate Rab10 phosphorylation ∼2-fold (Fig 3F). We also performed the secondary screen monitoring LRRK2-mediated Rab12^Ser106^ phosphorylation instead of Rab10^Thr73^ phosphorylation (Fig 3A, C). This analysis revealed that most activating variants also enhanced Rab12 phosphorylation in a LRRK2 kinase dependent manner at Ser106. However, we observed three variants (A1442P (ROC), T2031S (kinase) and N2081D (kinase) that enhanced Rab10 but not Rab12 phosphorylation (Fig 3A, 3B, 3C). Thus, from the group of 100 LRRK2 variants analysed, we conclude that 23 activated (based on Rab10 phosphorylation), 75 had no effect and 2 variants [ROC (S1508R) and COR_A_ (A1589S)] significantly reduced LRRK2 kinase activity.

**Figure 2.**
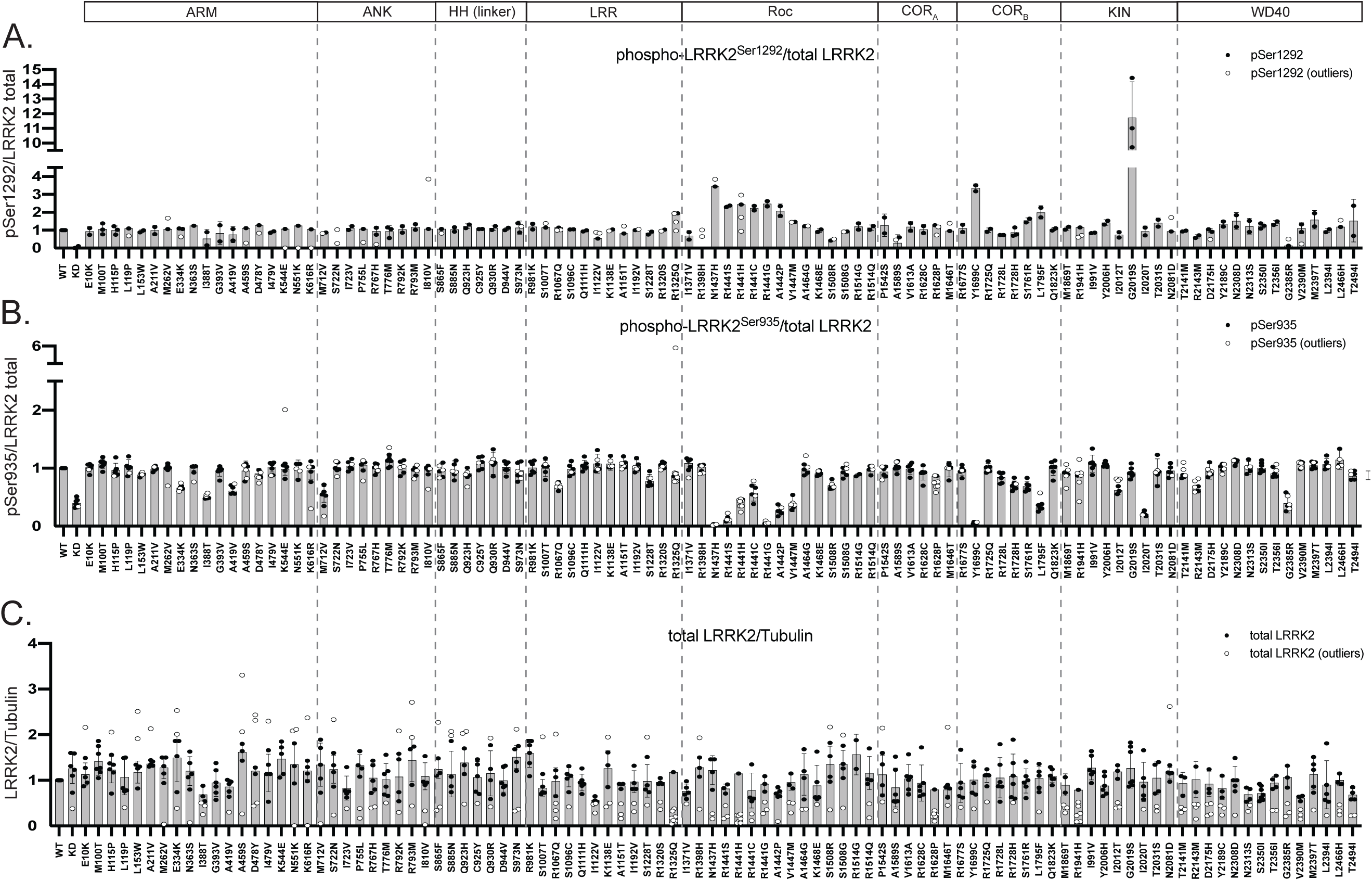
Quantitative analysis of phosphorylation and expression of selected PD and CD-associated LRRK2 variants assessed in primary screens. FLAG-tagged LRRK2 wildtype, kinase dead (KD = D2017A), and the indicated variants were transiently expressed in HEK293 cells. 24 hours post-transfection, cells were lysed and analysed by quantitative immunoblotting (as in Supplementary Figure 1). Quantified immunoblotting data are presented as ratios of phospho-LRRK2 Ser1292/total LRRK2 (A), phospho-LRRK2 Ser935 (B), and total LRRK2/Tubulin (C), normalized to the average of LRRK2 wildtype values for each replicate (mean ± SD). Combined immunoblotting data from 6 independent biological replicates are shown. Dashed lines segment the graphs into corresponding regions of LRRK2 as listed in the domain schematic.

**Figure 3.**
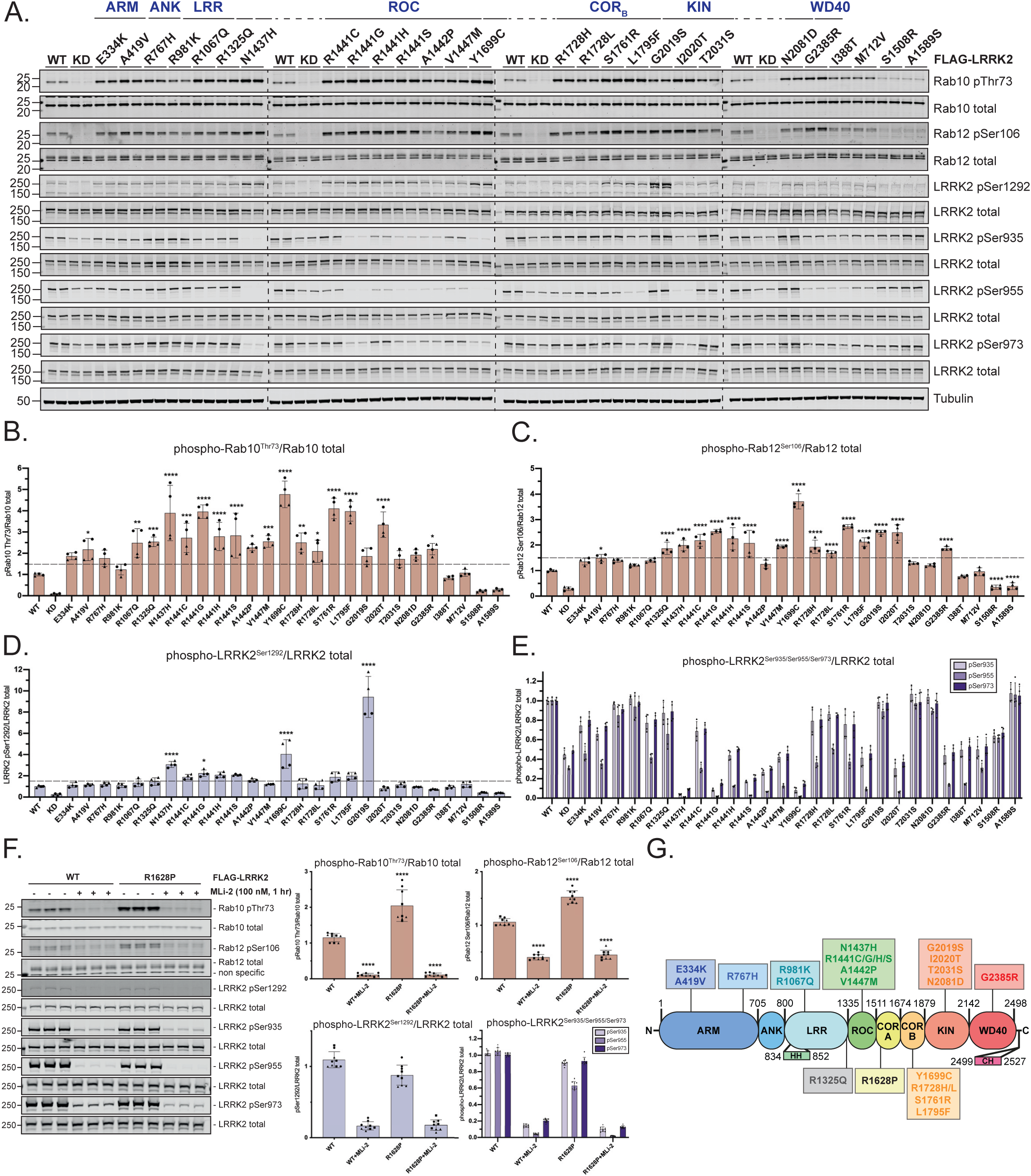
23 LRRK2 variants with mutations spanning multiple domains significantly augment LRRK2-mediated Rab10^Thr73^ phosphorylation. (A) FLAG-tagged LRRK2 wildtype, kinase dead (KD = D2017A) and the indicated variants were transiently expressed in HEK293 cells. 24 hours post-transfection, cells were lysed and analysed by quantitative immunoblotting using the indicated antibodies. Each lane represents a different dish of cells. Data quantification is shown in panels B-E. (B-E) Quantified immunoblotting data are presented as ratios of pRab10 Thr73/total Rab10 (B), pRab12 Ser106/total Rab12 (C), phospho-LRRK2 Ser1292/total LRRK2 (D), phospho-LRRK2 Ser935/total LRRK2, phospho-LRRK2 Ser955/total LRRK2, or phospho-LRRK2 Ser973/total LRRK2 (E), normalized to the average of LRRK2 wildtype values for each replicate (mean ± SD). Combined immunoblotting data from 2 independent biological replicates (each performed in duplicate) are shown. Data were analysed using one-way ANOVA with Dunnett’s multiple comparisons test. Statistical significance was determined from four replicate values for each variant, and represented with p-values (*p < 0.05, ** p < 0.01, *** p < 0.001, **** p < 0.0001). (F) FLAG LRRK2 WT or R1628P was expressed in HEK293 cells. Each lane represents a different dish of cells. One hour prior to lysis, cells were treated with vehicle (0.1% v/v DMSO) or 100 nM MLi-2. Cell lysates were analysed by quantitative immunoblotting and quantified data are analysed and presented as in (A-E). Quantified data are representative of three independent experiments, each performed in triplicate. (G) Domain schematic of LRRK2 highlighting the position of the 23 LRRK2 variants selected for further analysis.

**Figure 4.**
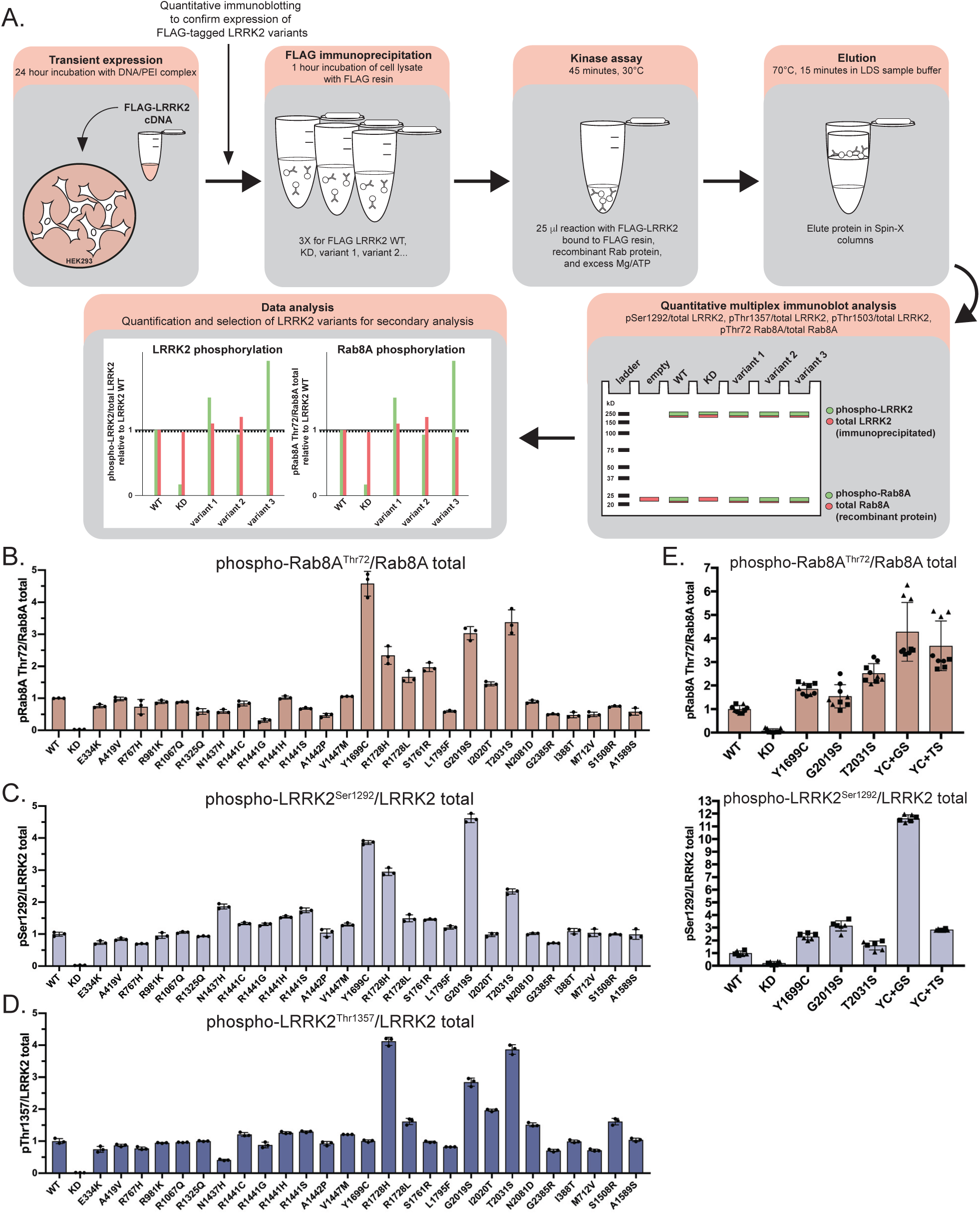
COR_B_ and kinase domain LRRK2 variants enhance *in vitro* LRRK2 kinase activity against recombinant Rab8A. (A) Workflow schematic outlining the immunoprecipitation kinase assay method employed to assess *in vitro* kinase activity of LRRK2 variants against recombinant Rab8A. Kinase reaction products were analysed by quantitative immunoblotting (as in Supplementary Figure 6, Supplementary Figure 7 and Supplementary Figure 8). (B-D) Data obtained from quantitative immunoblotting analysis of FLAG-LRRK2 immunoprecipitation kinase reactions for the indicates variants are presented as ratios of phospho-Rab8A Thr72/total Rab8A (B), phospho-LRRK2 Ser1292/total LRRK2 (C), and phospho-LRRK2 Thr1357/total LRRK2 (D) normalised to the average of LRRK2 wildtype values (mean ± SD). (E) Data obtained from quantitative immunoblotting analysis of FLAG-LRRK2 immunoprecipitation kinase reactions for the indicates variants are presented as ratios of phospho-Rab8A Thr72/total Rab8A, phospho-LRRK2 Ser1292/total LRRK2, relative to the average of LRRK2 wildtype values (mean ± SD).

The G2019S mutation stimulated LRRK2 Ser1292 autophosphorylation ∼10-fold, to a greater extent than other variants (Fig 2A, Fig 3A, D, SFig 2). Nine variants [ROC (N1437H, R1441G/C/S/H, A1442P) and COR_B_ (Y1699C, S1761R, L1795F)], increased Ser1292 autophosphorylation 2- to 4-fold (Fig 2A, Fig 3A, D, SFig 2).

Previous work revealed that variants that stimulate LRRK2 kinase activity, such as ROC (R1441G/C) and COR_B_ (Y1699C), suppressed LRRK2 biomarker phosphorylation, likely by promoting the closed, active conformation of the LRRK2 kinase domain [28–30]. Consistent with this, 10 activating variants [ROC (N1437H, R1441G/H/S, A1442P, V1447M), COR_B_ (Y1699C, L1795F), kinase (I2020T) and WD40 (G2385R)], displayed >2-fold reduction in phosphorylation of all biomarker sites (Fig 2B, Fig 3A, E, SFig 3). Seven variants [(ARM (E334K and A419V), LRR (R1067Q), ROC (R1441C) and COR_B_ (R1728H/L and S1761R)] showed reduced Ser955 phosphorylation, with a moderate impact on Ser935 and Ser973 phosphorylation (Fig 2B, Fig 3A, E, SFig 2, SFig 3). The reduced activity variants [ROC (S1508R) and COR_A_ (A1589S)], possessed similar biomarker phosphorylation as wildtype LRRK2 (Fig 2B, Fig 3A, F, SFig 2, SFig 3). Two variants [ARM (I388T) and ANK (M712V)] decreased biomarker site phosphorylation without impacting LRRK2-dependent Rab10^Thr73^ phosphorylation (Fig 1C, Fig 2B, Fig 3A, F, SFig 2, SFig 3). Although the majority of the 23 variants that stimulate LRRK2 activity reduce biomarker site phosphorylation, variants located within the ANK (R767H), LRR (R1325Q) or kinase (G2019S, T2031S and N2081D) domains do not reduce Ser935 or other biomarker sites. None of the variants studied increased the basal level of phosphorylation of the biomarker sites.

We observed that the R1398H protective variant [51] displayed similar pRab10^Thr73^, pLRRK2^Ser1292^ and LRRK2 biomarker phosphorylation levels compared to that of wildtype LRRK2 (Fig 1C, Fig 2, SFig 2, SFig 3). We also analysed the effect of the R1398H mutation on the kinase activity of three different LRRK2 pathogenic variants (R1441G, Y1699C and G2019S) and found that this mutation does not significantly reduce LRRK2 activity within these variants (SFig 5). Most variants analysed did not markedly impact LRRK2 expression levels in HEK293 cells (Fig 2C, SFig 2).

**Figure 5.**
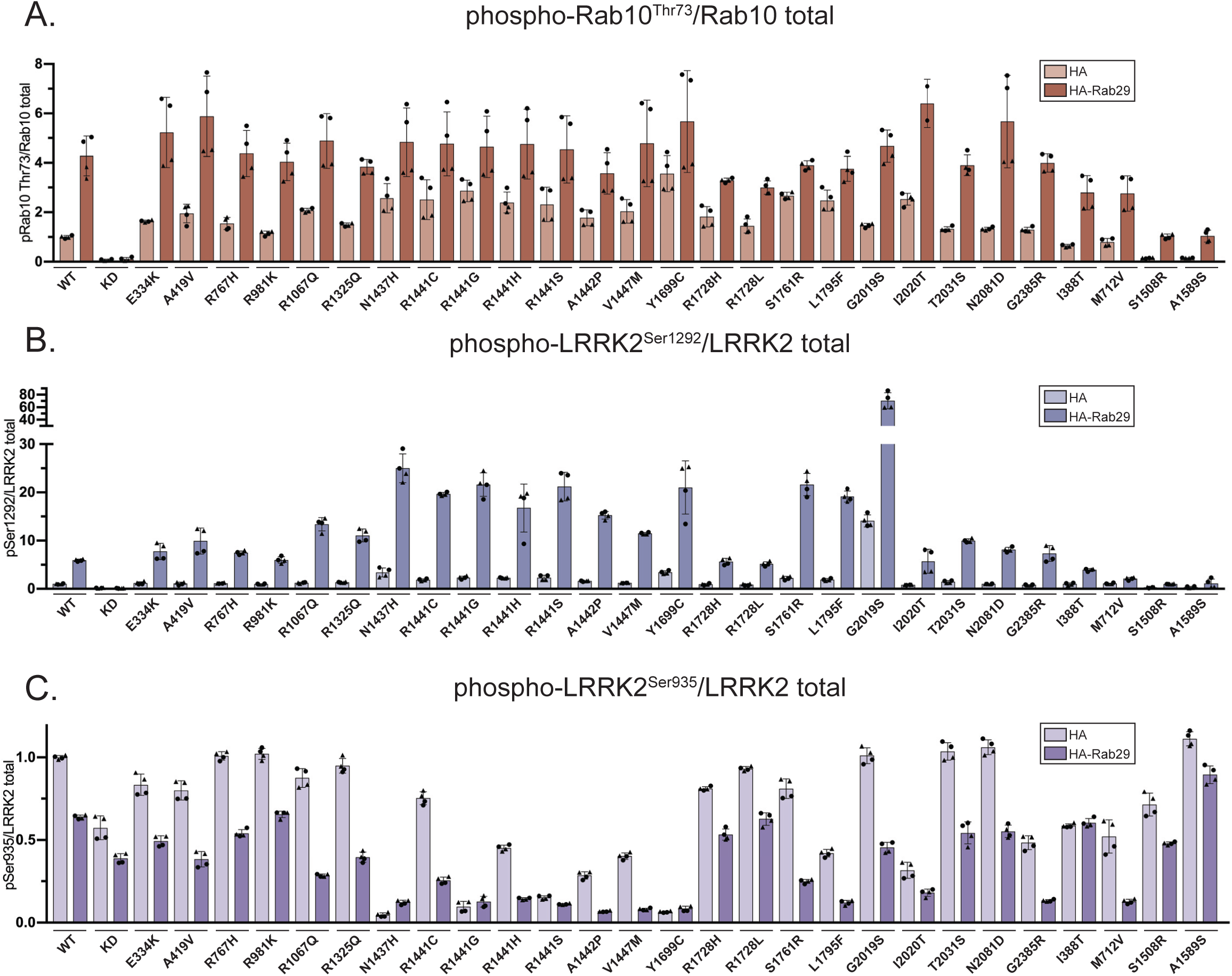
Selected LRRK2 variants are activated by Rab29. FLAG-tagged LRRK2 wildtype, kinase dead (KD = D2017A) and the indicated variants were transiently expressed in HEK293 cells with HA empty vector or HA-Rab29. 24 hours post-transfection, cells were lysed and analysed by quantitative immunoblotting (as in Supplementary Figure 9). (A-C) Quantified immunoblotting data are presented as ratios of phospho-Rab10/total Rab10 (A), phospho-LRRK2 Ser1292/total LRRK2 (B), phospho-LRRK2 Ser935/total LRRK2 (C), normalized to the average of LRRK2 wildtype values for each replicate (mean ± SD). Combined immunoblotting data from 2 independent biological replicates (each performed in duplicate) are shown.

### Impact of variants in an immunoprecipitation *in vitro* assay

Previous work revealed that pathogenic LRRK2 variants, such as the common G2019S kinase domain variant, directly enhance LRRK2 kinase activity, and this effect was recapitulated in recombinant *in vitro* kinase assays [52, 53]. In contrast, other pathogenic variants, such as ROC (R1441G), despite enhancing LRRK2 kinase pathway activity to a greater extent than the G2019S variant *in vivo*, failed to stimulate kinase activity of recombinant LRRK2 *in vitro* [53]. The contrasting effects on *in vitro* kinase activity suggest that these variants activate LRRK2 in cells by a different mechanism. This prompted us to investigate which of the *in vivo* activating variants enhanced activity of recombinant LRRK2 in an *in vitro* kinase assay. We expressed and immunopurified FLAG-tagged wildtype or mutant LRRK2 in HEK293 cells and subjected the purified protein to an *in vitro* kinase assay employing recombinant Rab8A as a substrate (Fig 4A). LRRK2-mediated phosphorylation of Rab8A at Thr72 was quantified using a previously characterized pan-selective phospho-Rab antibody [44](Fig 4A). We observed that of 22 variants which were found to enhance LRRK2 kinase activity as measured by pRab10^Thr73^ in the cellular assay, only 7 enhanced Rab8A phosphorylation *in vitro* by >1.5-fold relative to wildtype LRRK2 [COR_B_ (Y1699C, R1728H, R1728L, S1761R) and kinase (G2019S, I2020T and T2031S)] (Fig 4B, SFig 6, SFig 7). The kinase G2019S variant enhanced *in vitro* Rab8A phosphorylation around 3-fold, while the COR_B_ Y1699C and kinase T2031S variants stimulated activity ∼4-fold (Fig 4B, SFig 6, SFig 7). None of the variants within the ARM, ANK, LRR, ROC or WD40 domains enhanced immunoprecipitated LRRK2 activity *in vitro* (Fig 4B, SFig 6). The variants displaying reduced activity in the cellular assay [ROC (S1508R) and COR_A_ (A1589S)] possessed similar *in vitro* kinase activity towards Rab8A as the immunoprecipitated wildtype LRRK2 (Fig 4B, SFig 6), suggesting that these may impact LRRK2 kinase pathway activity in cells by an indirect mechanism rather than having a direct effect on LRRK2 kinase activity. Four variants [COR_B_ (Y1699C, R1728H) and kinase (G2019S, T2031S)] enhanced Ser1292 autophosphorylation over 2-fold in an *in vitro* assay (Fig 4C, SFig 6, SFig 7). We also studied autophosphorylation of LRRK2 at Thr1357 [54] and Thr1503 [55] employing recently developed phospho-antibodies. This revealed that the COR_B_ (R1728H), as well as the 3 kinase variants (G2019S, I2020T and T2031S), enhanced autophosphorylation of these sites ∼2-to 4-fold (Fig 4D, SFig 6, SFig 7). Since both COR_B_ and kinase domain variants increase LRRK2 activity in an immunoprecipitation assay, we next explored the impact of combining COR_B_ and kinase domain activating variants on LRRK2 kinase activity *in vitro*. This revealed that the Y1699C+T2031S as well as the Y1699C+G2019S combination increased LRRK2-mediated Rab8A phosphorylation to a greater degree than individual mutations assayed in parallel experiments (Fig 4E, SFig 8B). The Y1699C+G2019S combination also increased Ser1292 autophosphorylation *in vitro*, ∼12-fold, to a significantly greater extent than any other combination of mutations tested (Fig 4F, SFig 8B).

**Figure 6.**
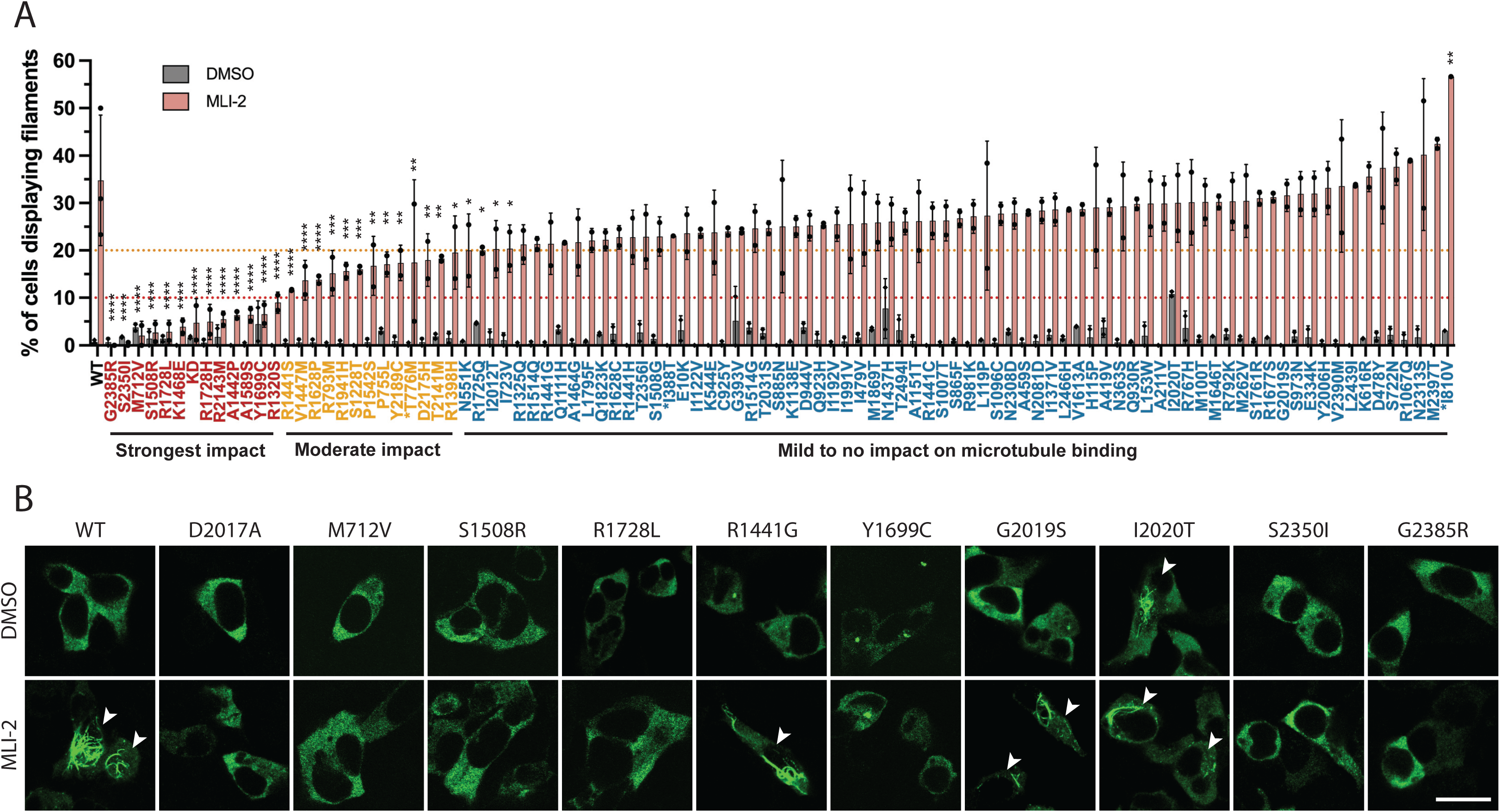
Impact of 98 LRRK2 variants on Type I inhibitor-induced microtubule association. (A) HEK293 cells transiently transfected with Flag-tagged LRRK2 wildtype, kinase dead (KD = D2017A) or the indicated variants were treated with 100 nM MLi-2 (or DMSO, control vehicle) for 3 hours to induce microtubule association. Cells were then fixed and subjected to immunofluorescent microscopy imaging of Flag-tagged LRRK2. Data are presented as % of LRRK2 signal-positive cells that show filamentous LRRK2. Bars represent mean ± SD and each circle represents a data point from an independent experiment with at least 50 Flag-LRRK2 staining-positive cells evaluated. The full experiment with 98 variants was performed twice and select few variants with lower expression levels were tested again separately in a third smaller scale experiment. Two-way ANOVA with the Dunnett’s multiple comparisons test was used to evaluate the statistical significance of the results (p values marked on the graph comparing the variant MLI-2-treated group to the WT MLi-2 treated group: * p<0.05, ** p<0.01, *** p<0.001, **** p<0.0001. None of the DMSO treated groups showed statistically significant differences from the WT group). Data are arranged by % of cells with filamentous LRRK2 signal upon MLI-2 treatment (low to high). (B) Sample images of the Flag-LRRK2 staining of selected variants. Scale bar - 10 μm. Cells with filamentous LRRK2 are marked with white arrowheads.

**Figure 7.**
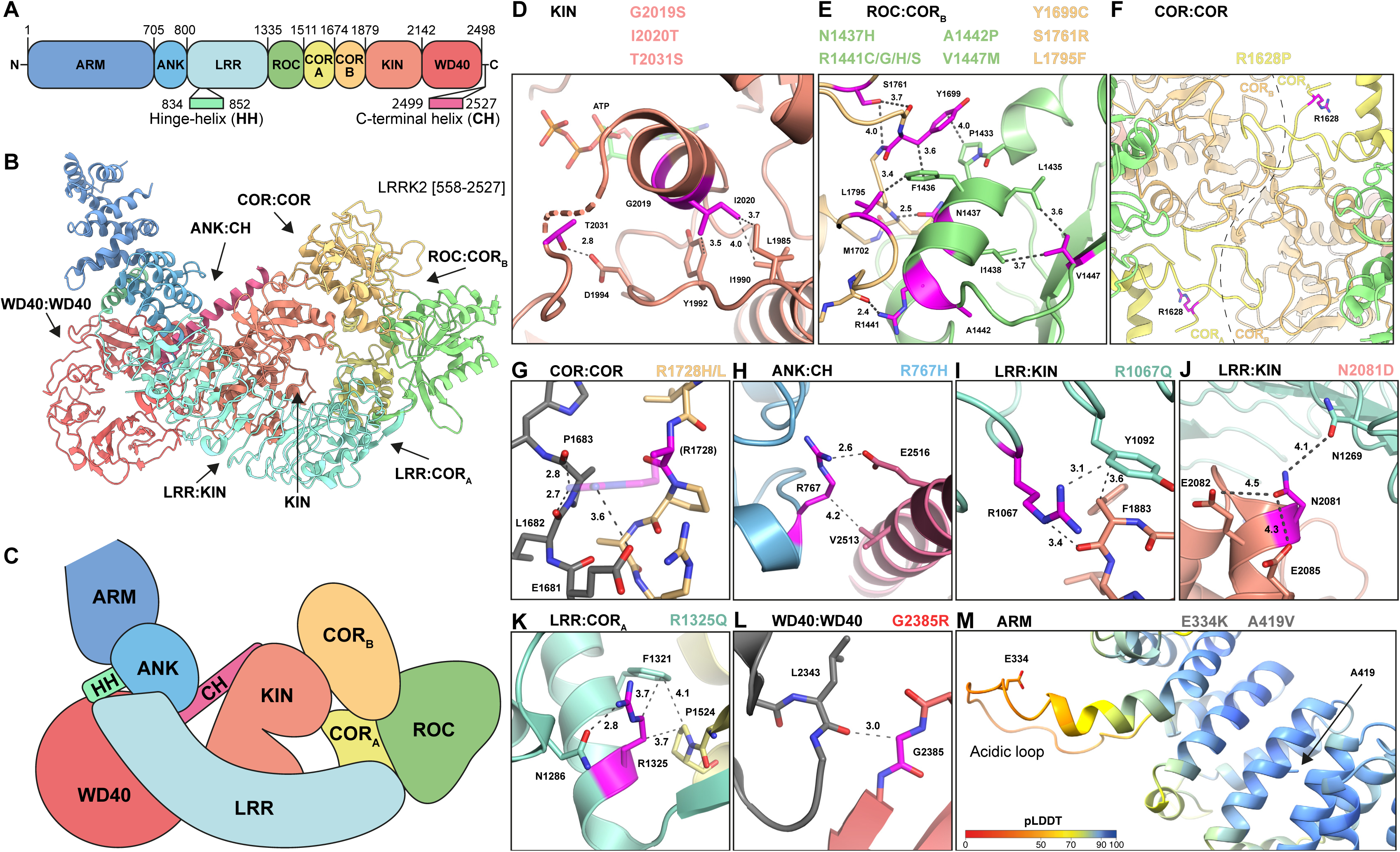
Structural analysis of identified activating LRRK2 variants. (A) Schematic domain overview of LRRK2 with domain boundaries. (B) Cartoon representation of LRRK2 [558–2527] with domains coloured as in A. Domain interfaces harbouring activating mutations and kinase domain are indicated by black arrows. (C) Schematic representation of LRRK2 domains as viewed in B. (D-L) Detailed views of LRRK2 variants in kinase active site (D), domain interfaces (E-L), colouring as in A and variants highlighted in magenta. Second LRRK2 molecule of dimer shown in grey (G, L). (G) R1728 side chain modelled in PyMOL shown as semi-transparent stick model. Distance measurements in Å are indicated by dark grey dashed lines. (M) Alphafold model of LRRK2 ARM domain coloured by local confidence score (pLDDT) with variant residues shown as stick models. LRRK2 structures used are PDB 7LI4 (B, D, E, H-K), PDB 7LHT (F, G), PDB 6DLO (L), AFDB AF-Q5S007-F1_v1 (M).

**Figure 8.**
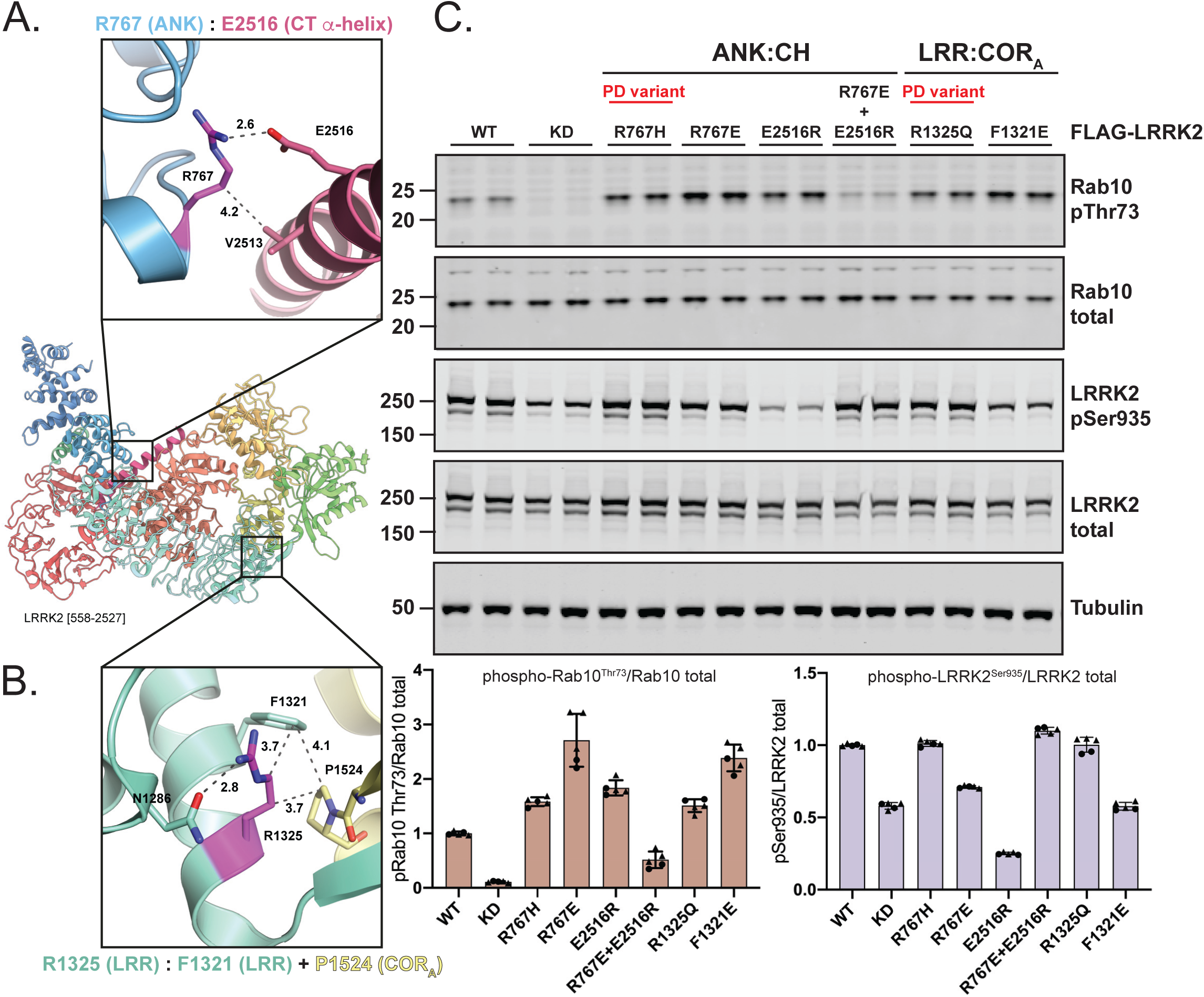
Structure guided mutations in the N-terminus of LRRK2 stimulate LRRK2-mediated Rab10 phosphorylation. (A-B) Cartoon representation of LRRK2 [558–2527] with detailed views of ANK (blue):CT α-helix (magenta) (A) and LRR (green):COR_A_ (yellow) (B) interactions. Distance measurements in Å are indicated by dark grey dashed lines. (Right panel) HEK293 cells were transfected with wildtype, kinase dead (KD = D2017A), and the indicated LRRK2 variants. Each lane represents a different dish of cells. Cells were harvested 24 hours post transfection and subjected to quantitative immunoblot analysis with the indicated antibodies. Each lane represents a different dish of cells. The ratios of phospho-Rab10 Thr73/total Rab10 and phospho-LRRK2 Ser935/total LRRK2 were normalized to wildtype LRRK2 values. Quantified data are presented as mean ± SD and are representative of two independent experiments.

### Activation of variants by overexpression of Rab29

Overexpression of Rab29 recruits LRRK2 to the Golgi membrane, promoting stimulation of LRRK2 kinase activity as assessed by increased Rab10^Thr73^ phosphorylation and LRRK2 Ser1292 autophosphorylation [11,12,56]. We next studied the impact of Rab29 overexpression on the activating variants and observed enhanced Rab10^Thr73^ phosphorylation and Ser1292 autophosphorylation with all variants tested (Fig 5, SFig 9). All ROC:COR_B_ domain interface variants enhanced Ser1292 autophosphorylation to a higher extent than the COR:COR interface variants (R1728H/L) following overexpression of Rab29 (Fig 5B, SFig 9). Rab29 also increased the activity of the two variants displaying reduced activity [ROC (S1508R) and COR_A_ (A1589S)] (Fig 5A). Co-expression of Rab29 decreased LRRK2 Ser935 phosphorylation of all variants (Fig 5C). All LRRK2 variants phosphorylated Rab29 at Thr71 to a similar extent (SFig 9), consistent with these being similarly activated by Rab29 binding.

**Figure 9.**
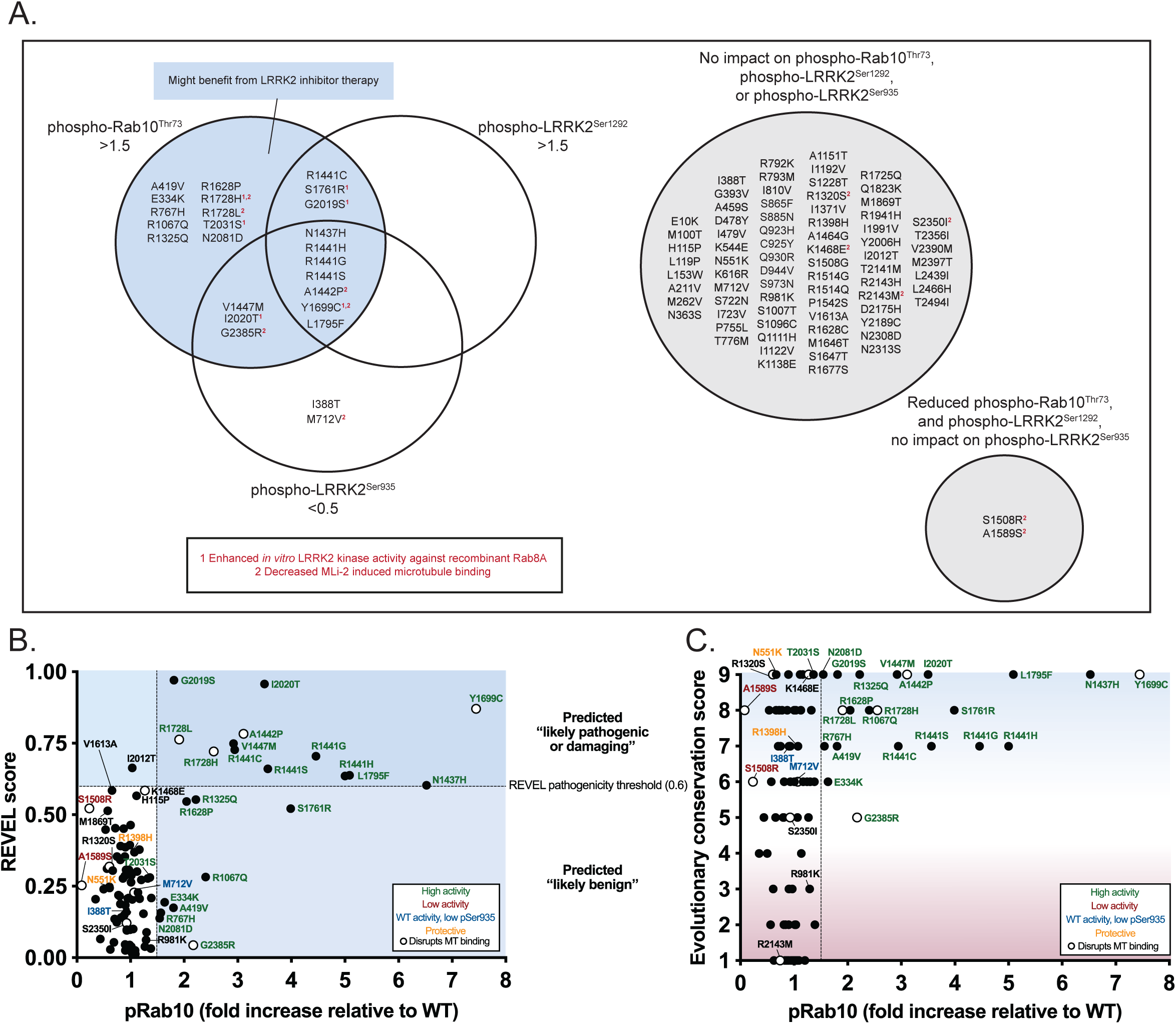
Correlation between activating LRRK2 mutations and high REVEL pathogenicity prediction or high evolutionary amino acid conservation scores. (A) Schematic summarizing biochemical data of 100 LRRK2 variants and categorisation of variants based on Rab10 phosphorylation, LRRK2 Ser1292 phosphorylation, and biomarker phosphorylation. Variants that enhance *in vitro* LRRK2 kinase activity or block MLi-2 induced microtubule binding are marked with a superscript highlighted in red. (B) REVEL scores for LRRK2 variants were acquired from Bryant et al 2021 or through the online pathogenicity prediction tool http://database.liulab.science/dbNSFP. REVEL scores were plotted against phospho-Rab10/total Rab10 ratios acquired for each LRRK2 variant that were normalized to wildtype. High activity variants (phospho-Rab10 > 1.5-fold relative to wildtype) are marked in green, low activity variants (similar to kinase inactive LRRK2) are marked in red, protective variants are marked in yellow, and variants that block MLi-2 induced microtubule binding are represented with an open circle. The REVEL pathogenicity threshold is marked with a dashed line (0.6). Above this line are variants predicted to be “likely pathogenic or damaging,” and variants below this line are predicted to be “likely benign.” (C) LRRK2 orthologue sequences were acquired from OrthoDB. Orthologue sequences were aligned using MAFFT. The multiple sequence alignment of LRRK2 orthologues was submitted to the ConSurf server to determine evolutionary conservation scores for LRRK2 amino acids (1 is low conservation and 9 is high conservation). Conservation scores were plotted against pRab10^Thr73^/total Rab10 ratios acquired for each LRRK2 variant that were normalized to wildtype. Activating variants (pRab10^Thr73^> 1.5-fold relative to wildtype) are marked in green, low activity variants (similar to kinase inactive LRRK2) are marked in red, protective variants are marked in yellow, and variants that block MLi-2 induced microtubule binding are represented with an open circle.

### Impact of variants on MLi-2 induced microtubule association

As mentioned in the introduction, Type I LRRK2 inhibitors including MLi-2, promote ordered oligomerization of LRRK2 on filaments [24–27]. We next investigated how 98 of the 100 LRRK2 variants impacted MLi-2 induced microtubule association. Cells expressing wildtype or LRRK2 variants were treated ± 100 nM MLi-2 for 3 hours, prior to fixation with 4% (w/v) paraformaldehyde. Immunofluorescence analysis was performed blinded and the fraction of cells displaying filamentous LRRK2 was quantified by studying 50-221 LRRK2 signal-positive cells in two separate experiments (Fig 6A, B). In the absence of MLi-2, typically < 5% cells of wildtype LRRK2 and most of the variants displayed filamentous LRRK2. For a few variants [ARM (G393V, A419V), ROC (N1437H) COR_B_ (R1725Q), kinase (I2020T) and WD40 (T2494I)], moderately elevated filamentous LRRK2 was observed in the absence of MLi-2. Consistent with previous work, MLi-2 treatment markedly increased the proportion of cells displaying filamentous LRRK2 to above 20% for wildtype and most studied variants. On the contrary, the kinase inactive LRRK2[D2017A] displayed no significant increase in filament formation following MLi-2 administration (Fig 6A, B). In addition, we observed that microtubule association in the presence of a Type I inhibitor was substantially reduced to below < 10% of cells for 12 variants [ARM (M712V), LRR (R1320S), ROC (A1442P, K1468E, S1508R), COR_A_ (A1589S), COR_B_ (Y1699C, R1728H/L) and WD40 (R2143M, S2350I, G2385R)] (Fig 6A, B). We also observed that a further 13 variants [ANK (P755L, T776M, R793M), LRR (S1228T), ROC (R1398H, R1441S, V1447M), COR_A_ (P1542S, R1628P), kinase (R1941H) and WD40 (T2141M, D2175H, Y2189C)] displayed a moderate reduction in MLi-2 induced microtubule association (Fig 6A).

### Predicted impact of activating variants on LRRK2 structure

The 23 identified activating variants are located across all domains of LRRK2 apart from the Hinge-helix and C-terminal helix (Fig 3A). Utilizing available high-resolution structures of inactive full-length LRRK2 and LRRK2 WD40 domain dimer (Protein Data Bank (PDB) 7LI4, 7LHT [8], PDB 6DLO [36]), and the LRRK2 model from the EMBL-EBI AlphaFold database (AFDB) [57] (Fig 7A, B, C), we analysed how these variants may impact LRRK2 structure and function. Three activating variants (G2019S, I2020T and T2031S) locate to the kinase active site (Fig 7D). Previous structural studies and molecular dynamics analyses of the G2019S and I2020T mutants suggest that they activate LRRK2 through changes in the flexibility of the kinase activation segment [8,58–60]. The T2031S variant is also located in the activation segment and may have a similar mechanism of kinase activation as G2019S and I2020T. An intriguing common feature of most of the other activating variants outside of the kinase domain is that they appear to destabilise LRRK2 interdomain interfaces. Most of these activating LRRK2 variants locate to the ROC and COR_B_ domains and lie within or nearby to the ROC:COR_B_ interface. These residues either participate in the interaction of the two domains directly [ROC (N1437H, R1441G/C/H/S), COR_B_ (Y1699C, L1795F)] or stabilize elements of the interface indirectly [ROC (A1442P, V1447M), COR_B_ (S1761R)] (Fig 7E). In addition, the COR_A_ variant R1628P and COR_B_ variants R1728H/L locate to the COR:COR dimer interface of LRRK2 (PDB 7LHT, Fig 7F, 7G). R1628 is located at the end of a loop in COR_A_ (residues 1613-1630) that interacts with the COR_B_ domain of the neighbouring LRRK2 molecule, and the R1628P mutation may place this loop in an unfavourable conformation, hindering dimerization (Fig 7F). While the R1728 side chain is not fully resolved in the LRRK2 dimer structure, it is likely to form hydrogen bonds with the carbonyl backbone of P1683 and L1682, as well as the side chain of E1681 (Fig 7G). The activating variants in the ROC and COR_B_ domains are therefore predicted to destabilize the ROC:COR_B_ and COR:COR interfaces. Further, the R767H variant likely destabilises the ANK:CH interface, as the arginine side chain of the R767 bridges the ANK domain to the C-terminal helix (CH) through hydrophobic and polar interactions with V2513 and E2516, respectively (Fig 7H). The LRR variant R1067Q variant may disrupt the LRR:kinase interface, as R1067 interacts with the kinase N-lobe through polar interactions with the carbonyl-backbone of F1883 (Fig 7I). The CD-associated N2081D variant is also found in the LRR:kinase interface and forms a hydrophilic interaction with residues of the LRR domain (Fig 7J), and this mutation has been proposed to disrupt these interactions [8]. The R1325Q variant resides at the LRR:COR_A_ interface, where the aliphatic part of the arginine side chain is involved in hydrophobic contacts with F1321 and P1524, while the guanidinium group is engaged with N1286 via hydrophilic interactions (Fig 7K). The common G2385R risk factor variant maps to the WD40:WD40 dimer interface, and mutation of this residue to arginine is likely to cause steric clashes with the neighbouring LRRK2 molecule and has been reported to block dimerization of this domain ([36], Fig 7L).

Noteworthy, the G2385R variant may also exert its pathogenic effect through coulomb repulsion with R841 of the Hinge-helix as previously proposed [8]. Finally, two of the identified activating variants, E334K and A419V, map to the N-terminal region of the ARM domain, which is absent from currently available high-resolution structures. However, the AlphaFold model of this region in LRRK2 (residues 159-511) has high local confidence scores (pLDDT) and agrees well with the experimentally determined cryo-EM map of LRRK2 (AFDB Q5S007-F1, EMD-23352,[8]) (SFig 10). In the AlphaFold model, E334 locates to an unstructured loop (residues 328-347) that protrudes from the armadillo repeats region (Fig 7M). Interestingly, this loop is highly acidic with 11 Asp/Glu out of 20 total residues and charge reversal through the E334K variant may therefore change the nature and function of this acidic loop. In contrast, the second ARM variant A419V is buried in the armadillo repeats and is not solvent exposed, and mutation to Val is likely to disturb the ARM structure (Fig 7M). Together, the activating ARM variants point to a third mechanism in regulating LRRK2 kinase activity, not yet explained by existing structures. We speculate that these mutations may affect substrate access, as the N-terminal ARM has previously been implicated in Rab substrate binding [13].

### Exploring mechanism of LRRK2 activation by disruption of interdomain interfaces

A common theme emerging in our analysis of the identified activating variants is that the apparent disruption of interdomain interfaces leads to activation of LRRK2. We therefore decided to explore this hypothesis by introducing structure-guided mutations in the ANK:CH and LRR:COR_A_ interfaces. As outlined above, the two N-terminal, PD-associated variants namely R767H (ANK, domain, Fig 8A) and R1325Q (LRR domain, Fig 8B), activate LRRK2.

Structural analysis indicates that the R767H mutation would disrupt an ionic interaction with E2516 residue located within the long C-terminal alpha-helix domain. Consistent with this, we find that the E2516R mutation (not a PD associated variant) activates LRRK2 (Fig 8C). Reinstating this ionic interaction by generating charge reversed, double R767E and E2516R mutations restores LRRK2 activity to that of wildtype enzyme (Fig 8C). Similarly, the R1325Q activating variant in the LRR domain is predicted to disrupt a hydrophobic interaction with F1321 (LRR) and P1524 (COR_A_) (Fig 8B). Consistent with this, disrupting this hydrophobic network by generating a F1231E mutation is sufficient to activate LRRK2 (Fig 8C). Together with the structural analysis of activating PD variants, these experiments strengthen the hypothesis that a disruption of LRRK2 interdomain interfaces can cause an increase in LRRK2 kinase activity.

## Discussion

In this study, we employed a tried and tested workflow to assess the activity of 100 LRRK2 variants that have previously been linked in the literature to PD (STable 1). The data obtained with respect to how each variant impacts LRRK2-mediated phosphorylation of Rab10 at Thr73, LRRK2 biomarker phosphorylation at Ser935, and LRRK2 auto-phosphorylation at Ser1292 are summarized in Figure 9A. Most importantly, we have characterised a group of 23 variants that reproducibly and robustly stimulate LRRK2-mediated Rab10^Thr73^ substrate phosphorylation >1.5-fold in the HEK293 cell overexpression system (Fig 1 to 3). We investigated how the computational pathogenicity REVEL score [41] of each variant (STable 1) correlates with experimentally measured variant activity and observed that 13 of the 23 activating variants possessed a REVEL score of > 0.6, which is considered pathogenic (Fig 9B). These include most of the previously characterised pathogenic mutations, as well as some of the novel variants identified in this study [ROC (A1442P, V1447M), COR_B_ (L1795F, R1728L)]. The activating variants located outside of the catalytic domains [ARM (E334K, A419V), ANK (R767H), LRR (R1067Q, R1325Q)] displayed REVEL scores of < 0.6 suggesting that this algorithm may be less effective at predicting activating variants lying outside the catalytic domains (Fig 9B). Only one of the variants tested, I2012T, possessed a REVEL score > 0.6 and did not stimulate LRRK2 in our assays (Fig 9B). Thus, 74 of the 75 variants studied, that did not enhance LRRK2 activity, displayed a REVEL score of < 0.6 (Fig 9B). We next explored how the evolutionary conservation score of each variant residue (STable 1) correlates with cellular kinase activity (Fig 9C). For this analysis, the evolutionary conservation was calculated using the ConSurf Server [46] and given a score of 1 to 9, with 1 being the least conserved, 4 being average and 9 being the most conserved. Functionally important residues that play an important role in controlling kinase activity would be expected to be highly evolutionarily conserved. Consistent with this, 21 of the 23 LRRK2 activating variants possess a high evolutionary conservation score of 7, 8, or 9 (Fig 9C). Only two activating variants, namely E334K (score 6) and G2385R (score 5), possess a lower score. Interestingly, the LRR R981K variant that was selected for our secondary screen and found not to significantly increase LRRK2 activity (Fig 3), possessed a low REVEL (0.062) and conservation score (3), consistent with these parameters having some utility in predicting pathogenicity.

Our data suggest that the impact of variants on LRRK2 kinase activity is better assessed employing a cellular assay measuring LRRK2-dependent pRab10^Thr73^ levels (Fig 1 to 3), rather than an immunoprecipitation *in vitro* kinase assay (Fig 4), as only 7 of the 22 variants analysed, both enhanced activity in the cellular assay and stimulated LRRK2 kinase activity *in vitro* [COR_B_ (Y1699C, R1728H, R1728L, S1761R) and kinase (G2019S, I2020T and T2031S)]. It is likely that the mutants that activate LRRK2 in the *in vitro* assay, stabilize the active conformation of the kinase domain. Further work is required to understand the mechanism by which most of the other identified variants stimulate LRRK2 kinase activity in cells. It is possible that these variants facilitate membrane interaction and/or association with other factors that enhance LRRK2-mediated phosphorylation of Rab10^Thr73^. In future work it would be interesting to explore whether the Cys residue in the Y1699C variant becomes oxidized or forms a disulphide bond and whether this could contribute to cellular and or *in vitro* activation observed with this variant. We also observed that combinations of Y1699C with G2019S or T2031S stimulated *in vitro* LRRK2 kinase activity to a greater extent than each individual variant alone (Fig 4E) and this finding may be useful in generating more active LRRK2 constructs for future functional and/or structural analysis. The finding that 3 variants, namely A1442P (ROC), T2031S (kinase) and N2081D (kinase), enhanced Rab10 phosphorylation without impacting Rab12 phosphorylation (Fig 3) indicates that certain variants could differentially impact phosphorylation of Rab proteins. It would be interesting to explore this further, and this could be done using a recently described multiplexed mass spectrometry assay [61].

The majority of the 23 variants that stimulate LRRK2 activity reduce biomarker site phosphorylation. This is consistent with previous work and has been interpreted to imply that active variants might adopt a conformation in which the biomarker sites are poorly phosphorylated or more efficiently dephosphorylated by the upstream kinase or phosphatase that acts on these sites [29, 53]. Consistent with this, Type I inhibitors that stabilize the active conformation of LRRK2 induce dephosphorylation of these biomarker sites [28]. Type II inhibitors that stabilize the inactive conformation of LRRK2 do not induce dephosphorylation of these residues [30]. However, our results emphasize that several activating variants located within the ANK (R767H) or kinase (G2019S, T2031S and N2081D) domains do not significantly reduce the phosphorylation of Ser935 or other biomarker sites (Fig 2B, 3). The reasons for this are currently not known. Perhaps these variants when expressed in cells retain a conformation distinct from the other activating variants. All activating variants within the ROC, COR_A_ or COR_B_ domain display reduced biomarker site phosphorylation. Our data also emphasize that autophosphorylation at Ser1292 is not directly correlated with LRRK2 kinase activity toward Rab substrates. LRRK2 G2019S shows by far the highest levels of Ser1292 phosphorylation among the pathogenic mutants, while its activity towards Rab10 is lower than other LRRK2 pathogenic mutants (Fig 1C, 2A, 3).

We also identify 12 variants [ARM (M712V), LRR (1320S), ROC (A1442P, K1468E, S1508R), COR_A_ (A1589S), COR_B_ (Y1699C, R1728H/L) and WD40 (R2143M, S2350I, G2385R)], in addition to kinase inactive LRRK2[D2017A], that significantly suppressed microtubule association in the presence of the MLi-2 Type I LRRK2 kinase inhibitor (Fig 6). Previous studies have established that the ROC-COR-kinase-WD40 domain fragment is sufficient to mediate oligomerization onto microtubules [9], and revealed that mutations impacting the WD40:WD40 interface [24] or the COR:COR interface [8], block microtubule association in cells. These conclusions are confirmed by a recent study that also highlights that disrupting the COR:COR interface by introducing the R1731L/D mutation, or the WD40:WD40 interface by introducing the S2343D mutation, markedly blocks microtubule association [10]. These findings likely account for why we observed that the COR_B_ (R1728H/R1728L), as well as the WD40 (R2143M, S2350I, G2385R) variants that likely affect the COR:COR and WD40:WD40 interfaces, inhibit microtubule association. Recent analysis identified a set of key basic residues located within the ROC domain that directly interact with acidic microtubule residues and this interaction is disrupted by a ROC (R1501W) variant linked to PD [10]. The ROC S1508R variant that we identified to block microtubule association is an internally buried residue in the ROC domain that is located adjacent to the basic microtubule-interacting patch and could affect positioning of the basic microtubule-binding patch.

K1468E is on the ROC domain surface pointing towards the microtubule surface. This residue is not part of the characterised microtubule-binding basic patch but located nearby and could also participate in microtubule binding. If the kinase inactivating D2017A variant impacts MLi-2 binding, this would account for why this mutation blocks MLi-2 mediated microtubule association. A previous study has also noted that a kinase-inactivating mutation distinct to that used in this study, namely K1906R, also blocked LRRK2 from association with microtubules [26]. The mechanism by which ARM (M712V), LRR (R1320S), ROC (A1442P) and COR_B_ (Y1699C) variants interfere with microtubule binding is currently not clear and requires further investigation.

It has been suggested that the ability of pathogenic variants to associate with microtubules may be linked to Parkinson’s disease [9]. The finding that 7 activating variants [ROC (A1442P), COR_B_ (Y1699C, R1728H/L) and WD40 (R2143M, S2350I, G2385R)] displayed reduced microtubule association, highlights that further analysis is required to explore a potential link between microtubule binding and PD. Interestingly, a recent study characterizing the R1731L and R1731D mutations that interfere with the COR:COR dimer interface, also observed that these mutations enhanced LRRK2 kinase activity [10], similar to what we have observed with the R1728H/L variants. The mechanism by which disruption of the COR:COR interface promotes kinase activation is currently unknown. It should be noted that pathogenic Flag-tagged LRRK2 mutants display minimal filament formation in the absence of MLi-2 (Fig 6), which is in contrast to previous reports employing GFP-tagged pathogenic LRRK2 mutants [24–26]. It would be worth investigating whether the known properties of certain GFP variants [62] to dimerize/oligomerize could account for the increased microtubule binding observed in other studies.

LRRK2 is a large protein with 2527 residues; over 1000 variants have been reported thus far and many people with PD likely carry additional rare LRRK2 variants of unknown clinical significance [41]. With next generation sequencing becoming more readily available for people with PD and disease modification with targeted treatments including LRRK2 kinase inhibitors entering clinical trials [63, 64], there is increased urgency to have a concrete set of functional parameters and experimental workflows available to help clinicians assess whether any given LRRK2 variant is likely to increase kinase activity and therefore a likely driver of the disease. We advocate the use of the experimental HEK293 cellular assay (Fig 1 to 3) to assess whether a variant enhances LRRK2 kinase activity by measuring LRRK2-dependent Rab10^Thr73^ phosphorylation, rather than the *in vitro* kinase assay (Fig 4) or solely relying on predictive methodology. Although the REVEL pathogenic score is useful, our data highlight that the REVEL score fails to identify most activating variants lying outside the GTPase and kinase domains (Fig 9B). A conservation evolutionary score of 7 to 9 would also indicate that the variant lies within a functionally critical position of the protein and most activating variants displayed conservation scores within this range. In future work, we are planning to experimentally assess additional LRRK2 variants linked to PD as they are reported. We are aiming to deposit all experimental data, REVEL and evolutionary conservation scores into a publicly accessible database (e.g., in collaboration with the Movement Disorder Society Genetic mutation database (www.MDSGene.org)) as soon as data become available. We hope that this information will help clinicians interpret LRRK2 variants in terms of their pathogenicity for genetic counselling, stratify people with LRRK2 variants and make a case to prioritize these individuals for LRRK2-targeting clinical trials, as well as stimulate further mechanistic and structural analysis to better understand how these variants enhance LRRK2 kinase activity. There is also a need to validate similar functional workflows for other PD-associated genes, including the lysosomal enzyme glucocerebrosidase (GBA1) [65].

## Materials and Methods

### Reagents

MLi-2 LRRK2 inhibitor was synthesized by Natalia Shpiro (University of Dundee). Human recombinant Rab8A (1-207, DU47363) used for the immunoprecipitation kinase assays was obtained from the MRC PPU Reagents and Services (https://mrcppureagents.dundee.ac.uk).

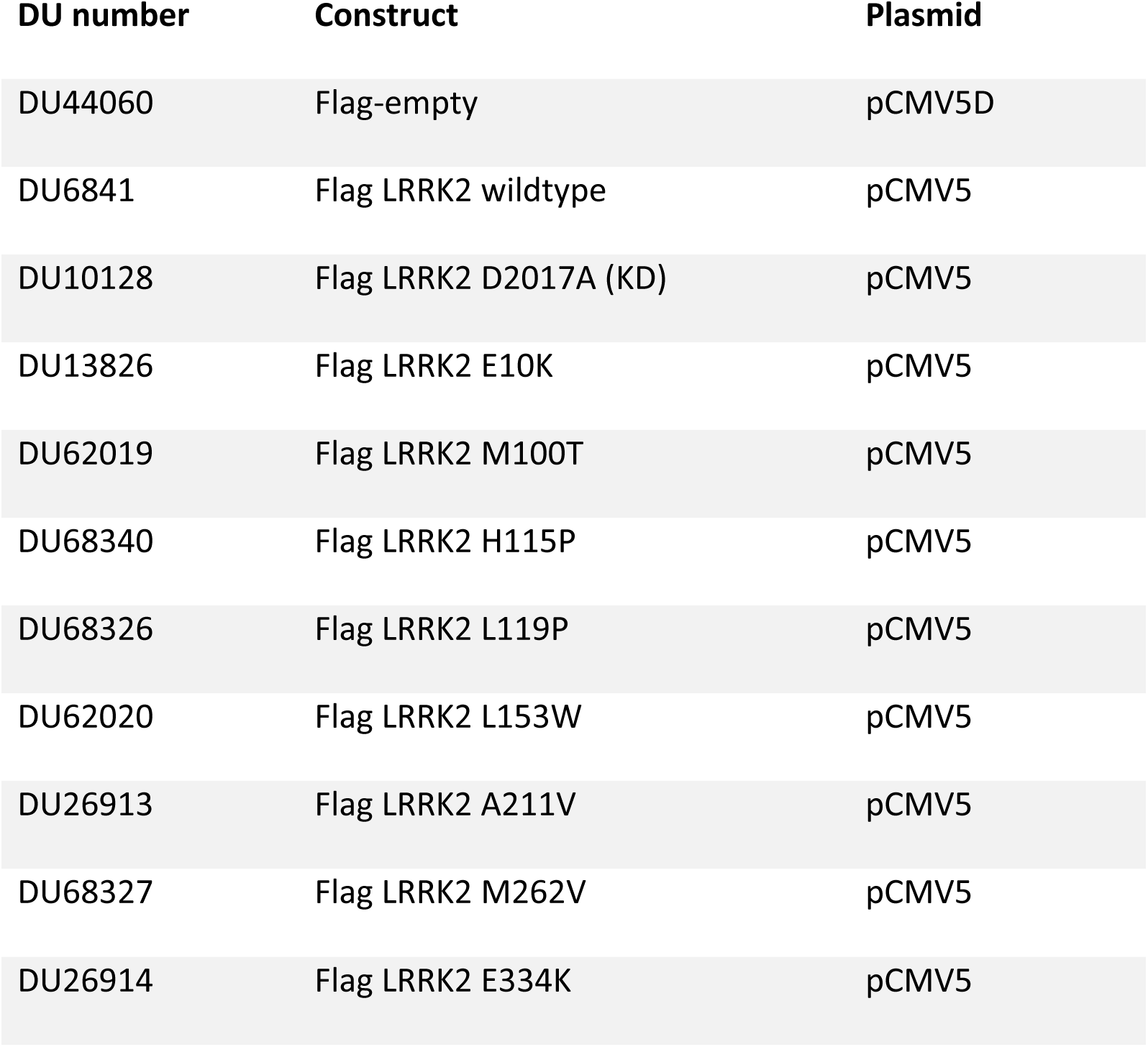
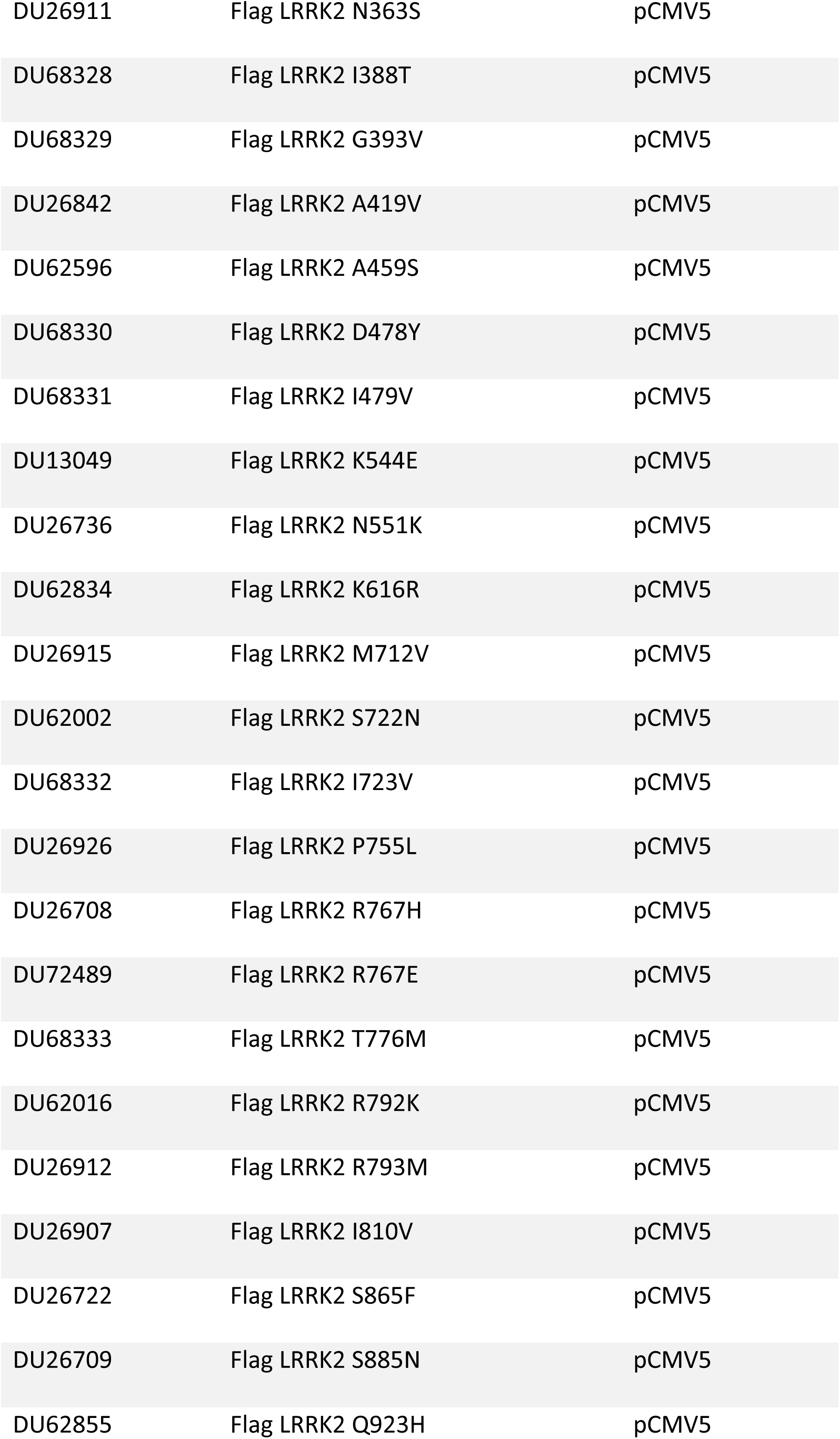
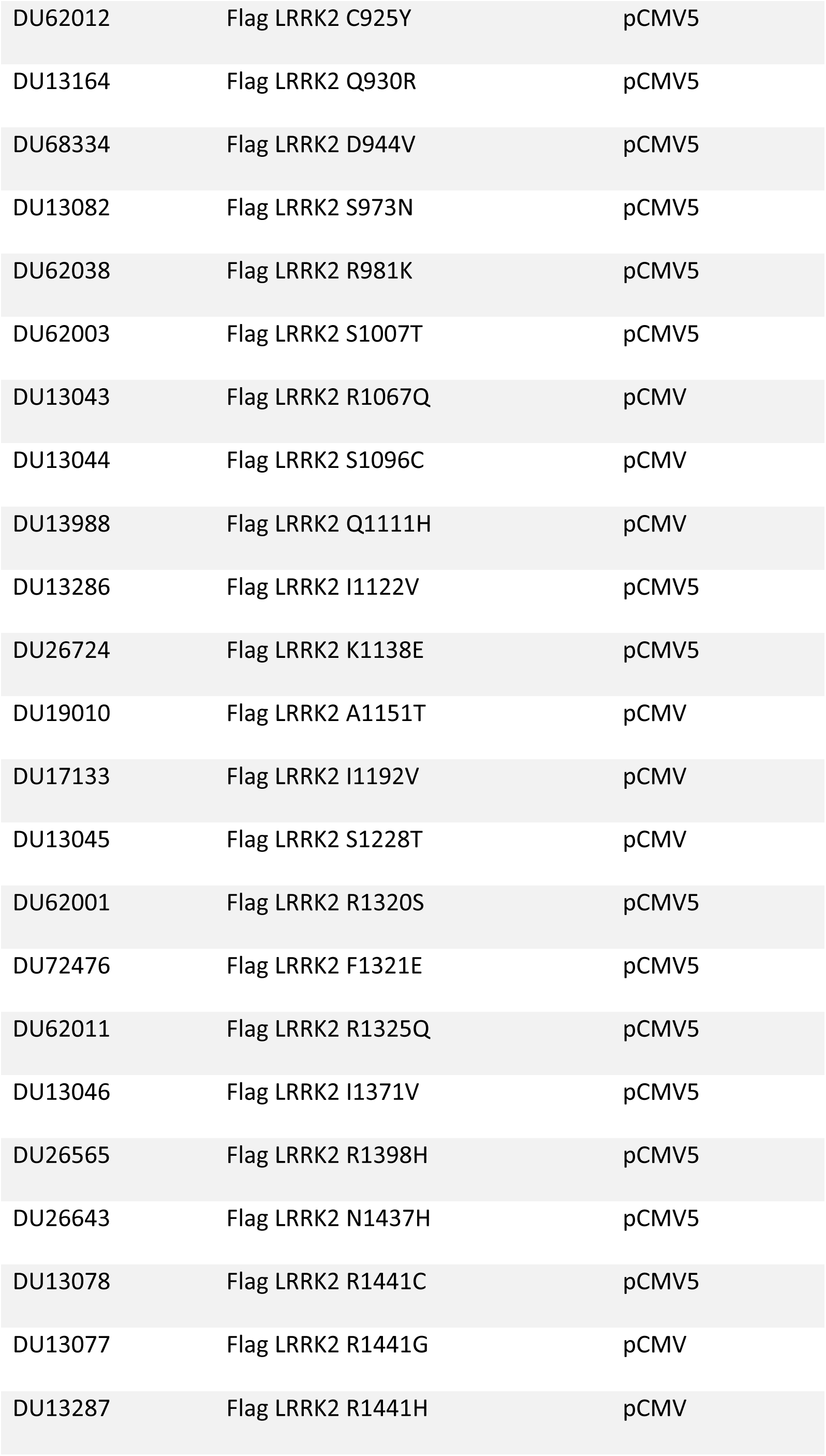
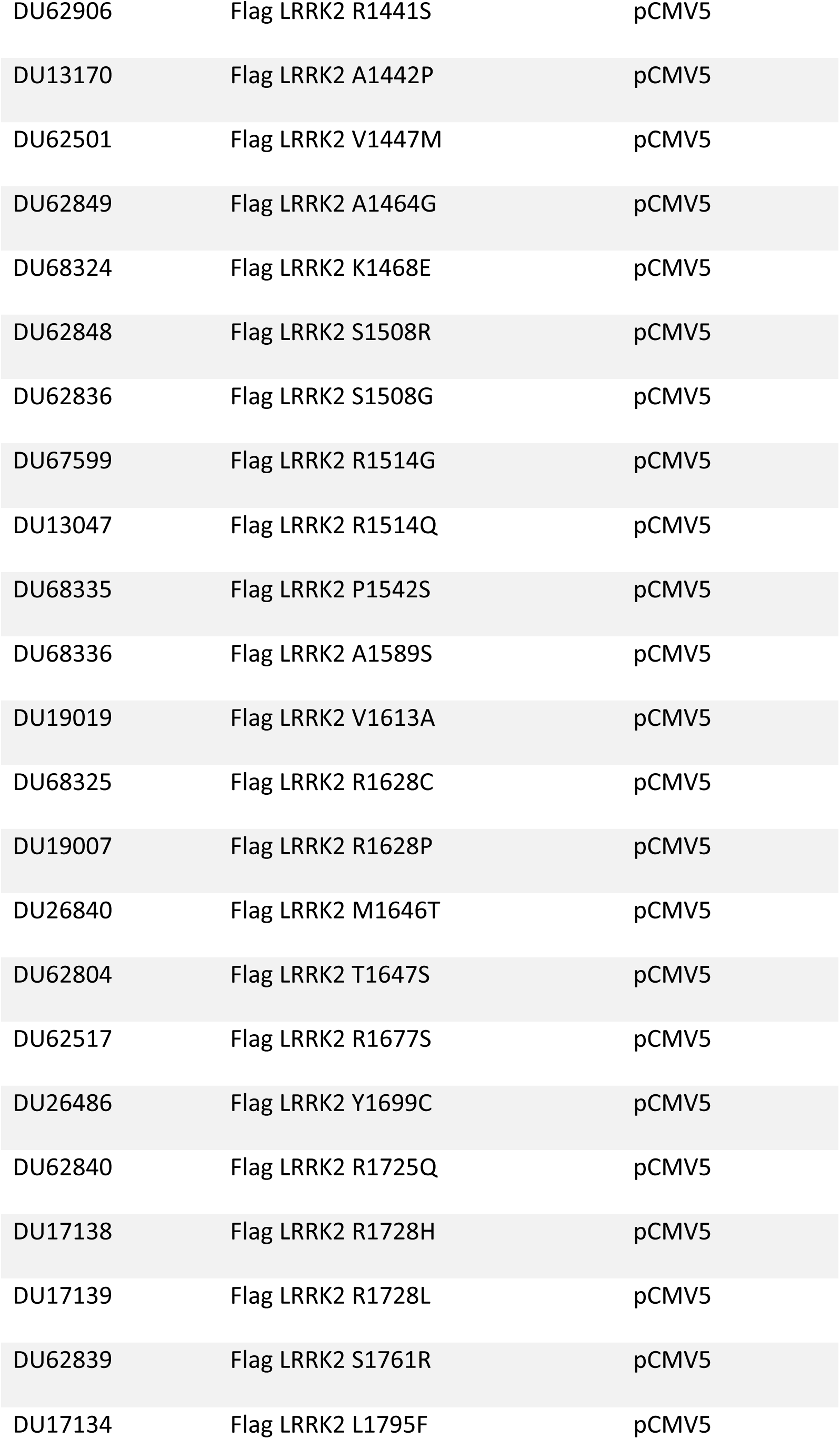
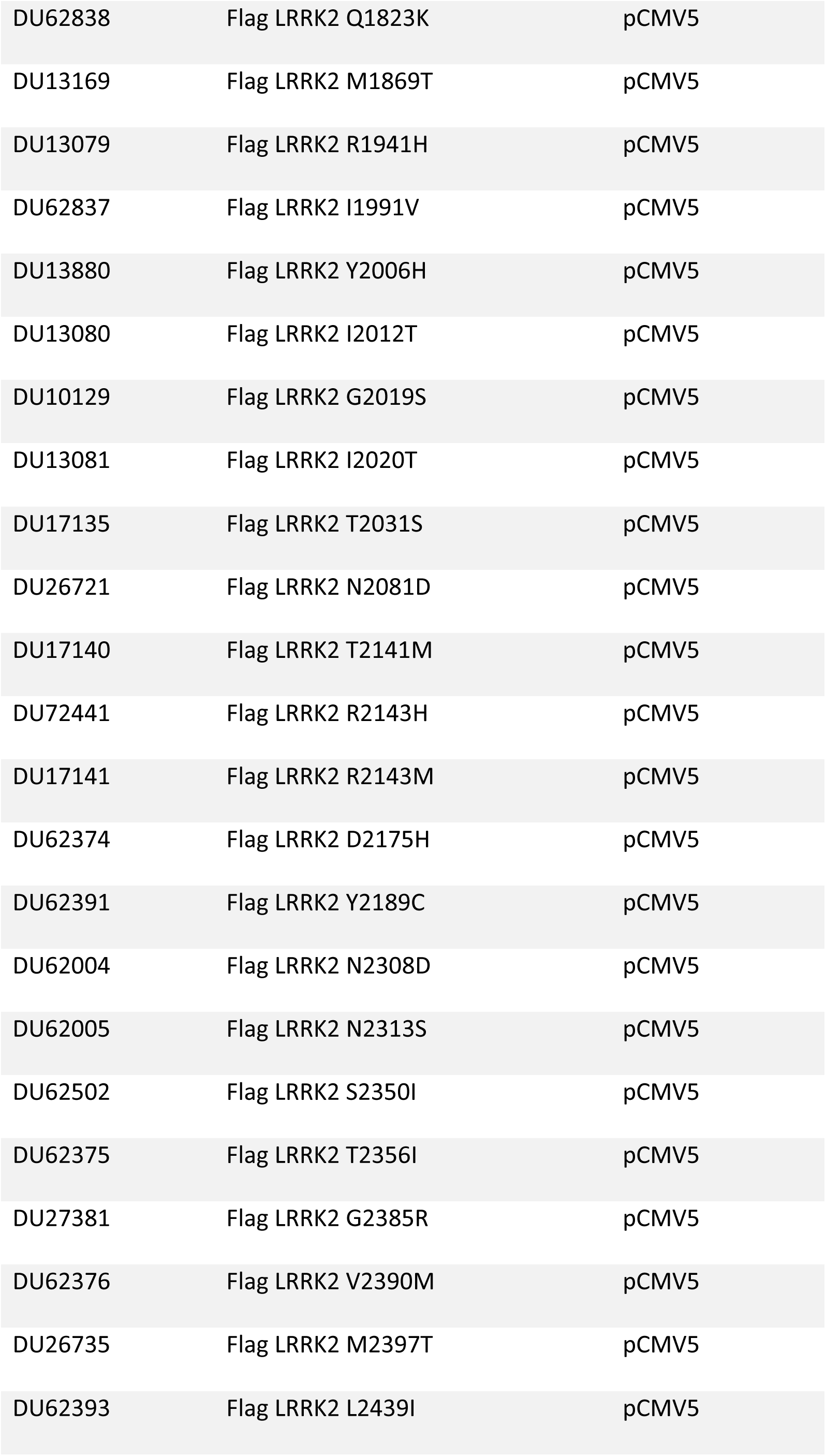
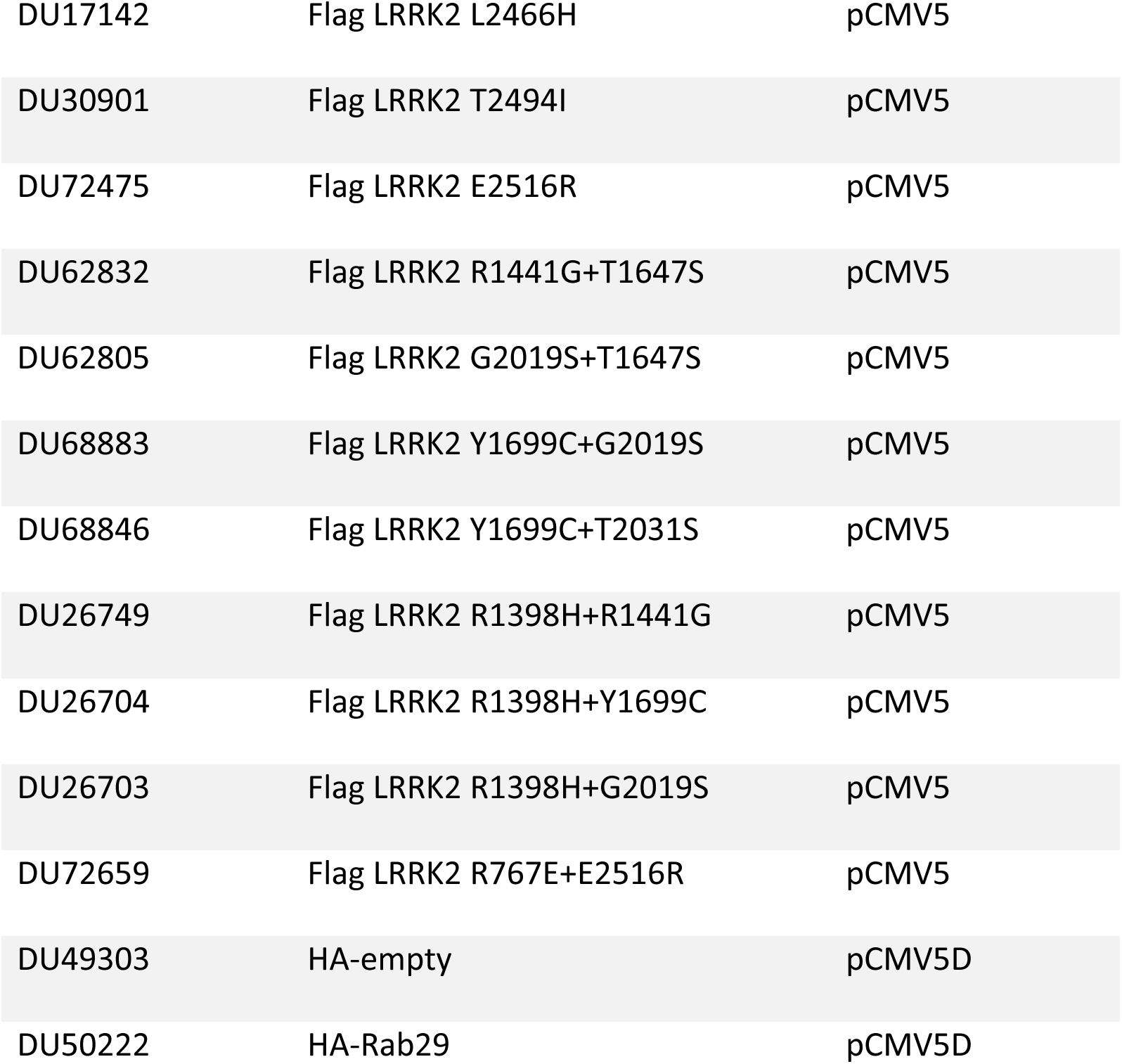

### Cell culture

HEK293 cells (ATCC Cat# CRL-1573, RRID:CVCL_0045) were cultured in DMEM (Dulbecco s Modified Eagle Medium) (Dulbecco’s Modified Eagle Medium, Gibco ™) containing 10% (v/v) foetal calf serum, 2 mM L-glutamine, 100 U/ml penicillin and 100 μg/ml streptomycin at 37°C in a humidified incubator maintaining 5% (v/v) CO2. Cells were regularly tested for mycoplasma contamination.

### Plasmids

All plasmids used in this study were obtained from the MRC PPU Reagents and Services (https://mrcppureagents.dundee.ac.uk). Each LRRK2 variant was confirmed by sequencing at the MRC Sequencing and Services (https://www.dnaseq.co.uk) and the amplified plasmid preparation quality was validated via agarose gel electrophoresis using ethidium bromide staining. All plasmids are available to request via the MRC PPU Reagents and Services website (https://mrcppureagents.dundee.ac.uk).

### Cell transfection and lysis

A protocols.io description of our cell transfection (dx.doi.org/10.17504/protocols.io.bw4bpgsn) and cell lysis method (dx.doi.org/10.17504/protocols.io.b5jhq4j6) has previously been described. For LRRK2 variant immunoblot analysis, HEK293 cells were seeded into 6-well plates and transiently transfected at 60-70% confluence using Polyethylenimine (PEI) transfection reagent with Flag-empty, Flag-LRRK2 wildtype or variant plasmids. 2 µg of plasmid and 6 µg of PEI were diluted in 0.5 mL of Opti-MEM™ Reduced serum medium (Gibco™) per single well. Cells were lysed 24 hours post-transfection in an ice-cold lysis buffer containing 50 mM Tris/HCl pH 7.4, 1 mM EGTA, 10 mM 2-glycerophosphate, 50 mM sodium fluoride, 5 mM sodium pyrophosphate, 270 mM sucrose, supplemented with 1 µg/ml microcystin-LR, 1 mM sodium orthovanadate, complete EDTA-free protease inhibitor cocktail (Roche), and 1% (v/v) Triton X-100. Lysates were clarified by centrifugation at 15000 *g* at 4°C for 15 min and supernatants were quantified by Bradford assay.

For LRRK2 variant screen for microtubule association, cells were split into either µ-Plate 24-wells (#1.5 polymer coverslip, black well, flat bottom, ibiTreat; Ibidi) for immunofluorescence or regular 24-well plates for immunoblotting control. Cells were transfected using Polyethylenimine (PEI) transfection with Flag-empty, Flag-LRRK2 wildtype or variant plasmids. 0.6 µg of plasmid and 1.7 µg of PEI in 0.15 mL of Opti-MEM™ per single well. Three hours prior to lysis, cells were treated with 100 nM MLi-2 or 0.1% (v/v) DMSO (vehicle). 48 hours post-transfection, cells for immunofluorescence were fixed for 10 minutes using 4% (v/v) PFA in PBS (Phosphate Buffered Saline), pre-warmed to 37°C and cells for immunoblotting were lysed as above.

### Quantitative immunoblot analysis

A protocols.io description of our quantitative immunoblotting protocol has previously been described (dx.doi.org/10.17504/protocols.io.bsgrnbv6). Briefly extracts were mixed with a quarter of a volume of 4x SDS-PAGE loading buffer [250 mM Tris–HCl, pH 6.8, 8% (w/v) SDS, 40% (v/v) glycerol, 0.02% (w/v) bromophenol blue and 5% (v/v) 2-mercaptoethanol] and heated at 95 °C for 5 minutes. Samples were loaded onto NuPAGE 4–12% Bis–Tris Midi Gels (Thermo Fisher Scientific, Cat# WG1402BOX or Cat# WG1403BOX) or self-cast 10% Bis-Tris gels and electrophoresed at 130 V for 2 hours with NuPAGE MOPS SDS running buffer (Thermo Fisher Scientific, Cat# NP0001-02). At the end of electrophoresis, proteins were electrophoretically transferred onto a nitrocellulose membrane (GE Healthcare, Amersham Protran Supported 0.45 µm NC) at 90 V for 90 minutes on ice in transfer buffer (48 mM Tris base and 39 mM glycine supplemented with 20% (v/v) methanol). The membranes were blocked with 5% (w/v) skim milk powder dissolved in TBS-T (50 mM Tris base, 150 mM sodium chloride (NaCl), 0.1% (v/v) Tween 20) at room temperature for 1 hour. Membranes were washed three times with TBS-T and were incubated in primary antibody overnight at 4 °C. Prior to secondary antibody incubation, membranes were washed three times for 15 minutes each with TBS-T. The membranes were incubated with secondary antibody for 1 hour at room temperature. Thereafter, membranes were washed with TBS-T three times with a 15 minute incubation for each wash, and protein bands were acquired *via* near infrared fluorescent detection using the Odyssey CLx imaging system and quantified using Image Studio Lite (Version 5.2.5, RRID:SCR_013715).

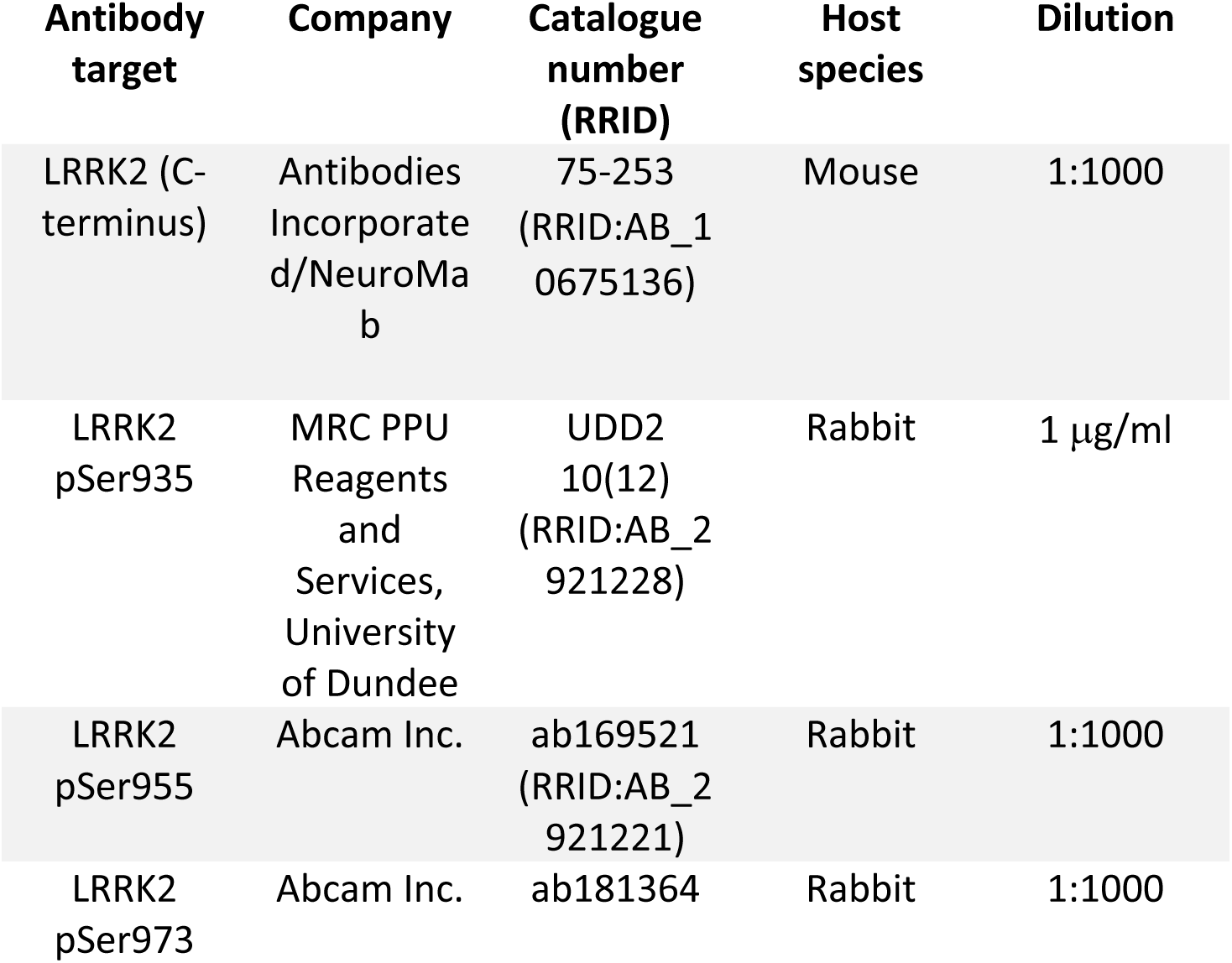
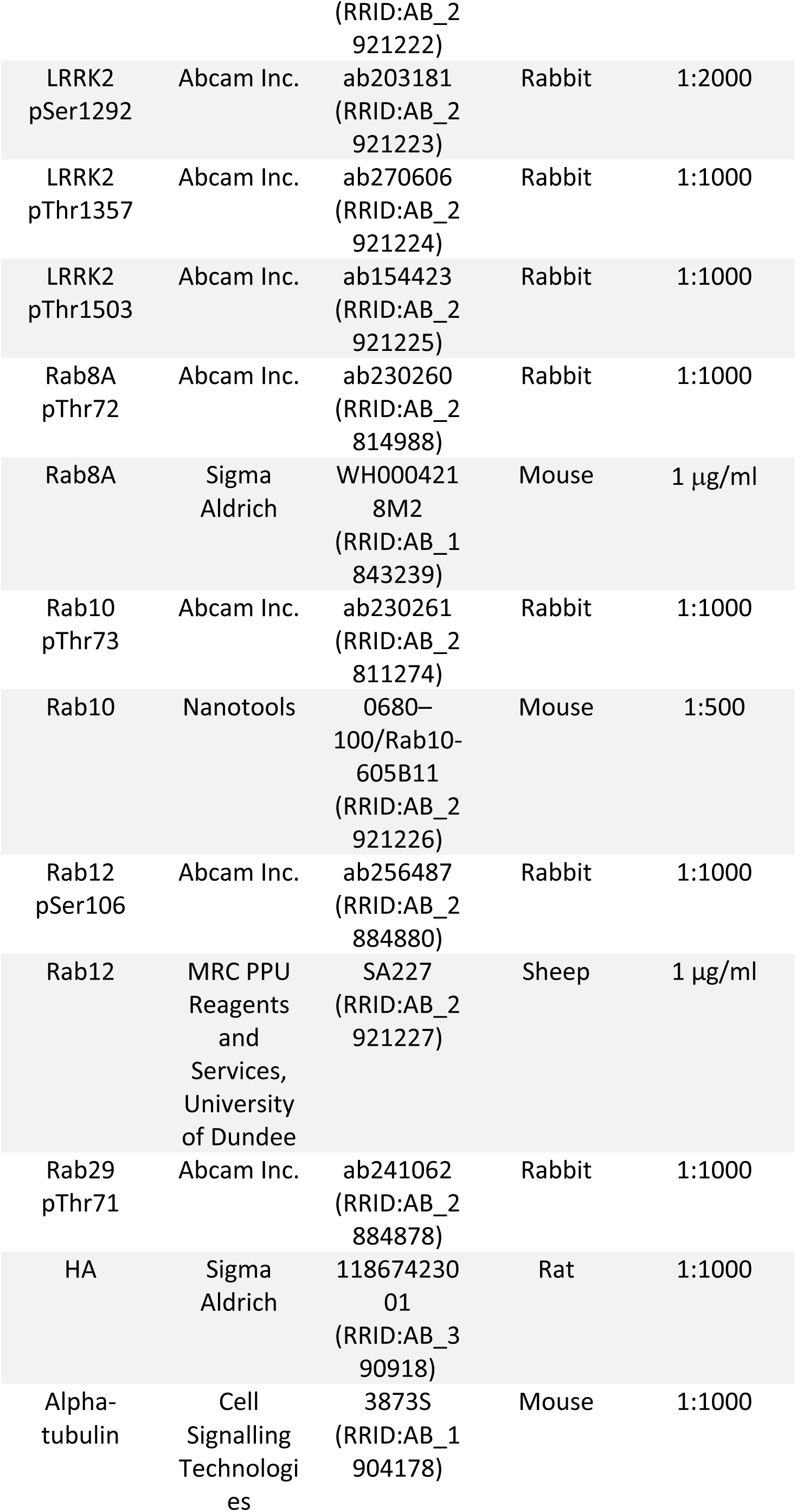
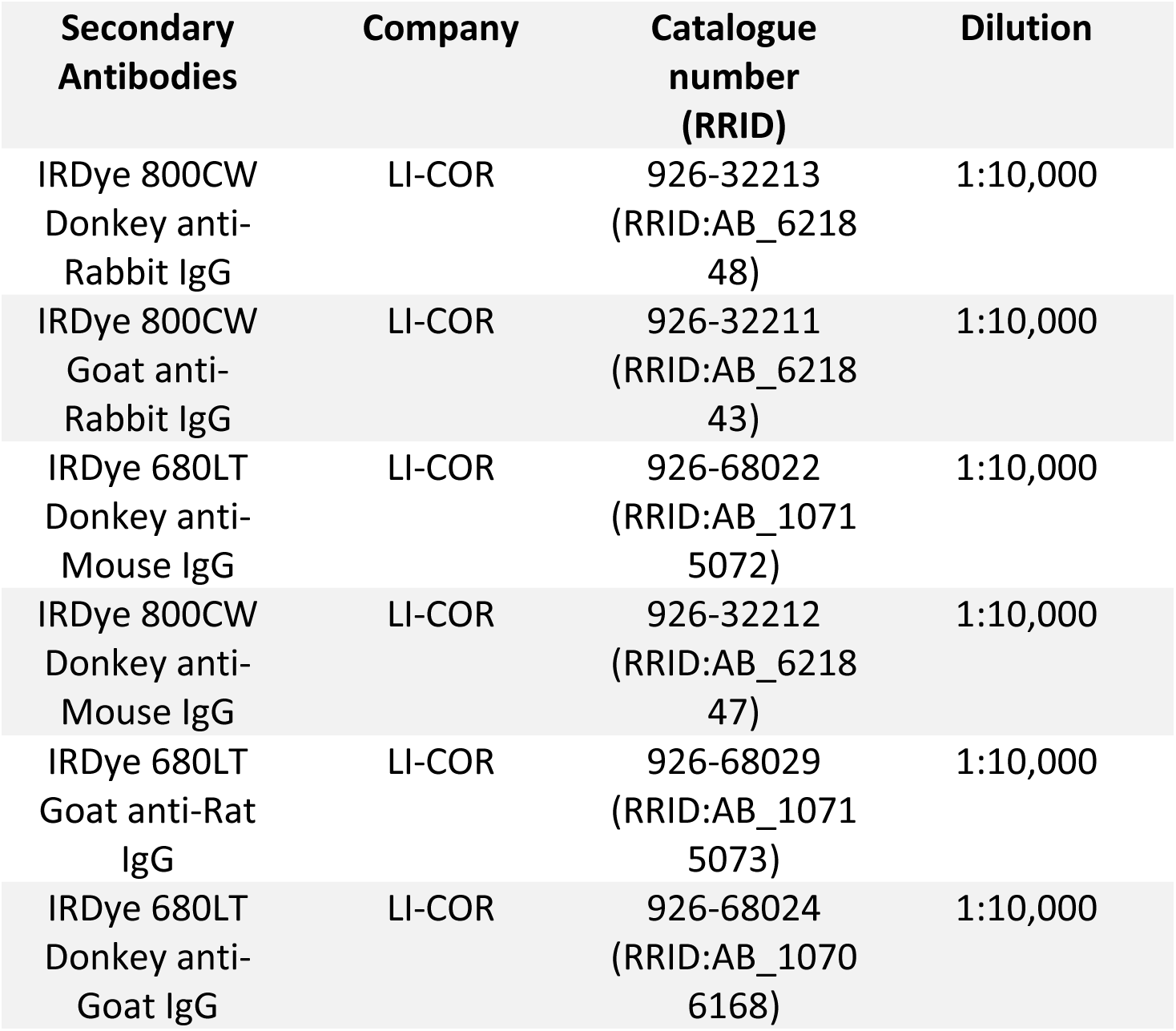

### Immunoprecipitation kinase assays

A protocols.io description of our LRRK2 immunoprecipitation kinase assay has previously been described (dx.doi.org/10.17504/protocols.io.bw4bpgsn). Briefly, HEK293 cells were transiently transfected with FLAG-LRRK2 wildtype, FLAG-LRRK2 D2017A, and FLAG-tagged LRRK2 variants using polyethylenimine (PEI) and lysed 24 hours post-transfection. Prior to immunoprecipitation, cell lysates were subjected to quantitative immunoblotting to assess the expression of each LRRK2 variant by quantifying total LRRK2 and normalizing to Tubulin. 100 µg cell lysate expressing FLAG-LRRK2 wild type, and the equivalent amount of cell lysate adjusted according to expression of each FLAG-tagged LRRK2 variant, was used to immunoprecipitate LRRK2 with 10 µl anti-FLAG M2 resin for one hour at 4°C, and immunoprecipitations were set up in triplicate per dish of cells. Immunoprecipitates were washed three times with lysis buffer supplemented with 300 mM NaCl, and twice with 50 mM Tris/HCl (pH 7.5). Kinase reactions were set up in a total volume of 25 µl, with immunoprecipitated LRRK2 in 50 mM Tris/HCl (pH 7.5), 10 mM MgCl2, 1 mM ATP, in the presence of 5 µg recombinant Rab8A. Kinase reactions were carried out at 30°C for 45 minutes at 1150 rpm. Reactions were terminated by adding 25 µl 4X LDS (lithium dodecyl sulfate) loading buffer to the beads. After heating the reactions at 70°C for 15 minutes, the eluates were collected by centrifugation through a 0.22 μM pore-size Spin-X column and supplemented with 2% (v/v) 2-mercaptoethanol. The kinase reactions were heated at 95°C for 5 minutes, then subjected to quantitative immunoblot analysis. Membranes were developed using the Licor Odyssey CLx scan Western Blot imaging system and quantified using Image Studio Lite (Version 5.2.5, RRID:SCR_013715).

### Immunofluorescence, imaging, and cell counting and quantitation of Microtubule binding

A protocols.io description of the Immunofluorescence-based method that we used to assess LRRK2 association with microtubules in HEK293 cells has been described (dx.doi.org/10.17504/protocols.io.b5jhq4j6). Cells were fixed in 4% (w/v) paraformaldehyde (Sigma Aldrich #P6148) in PBS, pH 7.4 for 10 minutes, followed by permeabilisation using 1% (v/v) NP-40 in PBS for 10 minutes. Cells were then blocked in 1% (w/v) bovine serum albumin in PBS for 1 hour at room temperature. Blocked cells were incubated with Flag M2 (raised in mouse, Sigma-Aldrich Cat# F1804, RRID:AB_262044, 1:1000 dilution) and β-tubulin (raised in rabbit, Abcam Cat# ab6046, RRID:AB_2210370, 1:500 dilution) antibodies for 2 hours at 37°C. Cells were washed 3 times (15 minutes each) with 0.2% (w/v) bovine serum albumin in PBS and incubated with secondary antibodies (goat-anti-mouse Alexa Fluor 488 Thermo Fisher Scientific Cat# A-21202, RRID:AB_141607 and goat-anti-rabbit Alexa Fluor 594 Thermo Fisher Scientific Cat# A-21207, RRID:AB_141637, 1:500 dilution) and 1 µg/ml DAPI (4’,6-Diamidino-2-Phenylindole, Dilactate) for 1 hour at room temperature in the dark. Cells were washed 3 times (15 minutes each) with 0.2% (w/v) bovine serum albumin in PBS and were kept in PBS at 4° C until imaging. Plates were imaged on the Zeiss LSM 710 or 880 laser scanning microscopes using the x40 EC Plan-Neofluar (NA 1.3) objective with a zoom of 0.6 and optical section thickness of 1.0 µm (image size 2048x2048 pixels, pixel size 0.173 mm). 4-6 randomly selected fields with Alexa Fluor 488(FLAG)-positive cells were collected for each well blinded to LRRK2 variant and treatment conditions. For further cell counting, LRRK2 variant and treatment condition were blinded from the counter by renaming the image files using a simple Python code script (IPython (RRID:SCR_001658). The Python code script used to rename image files for blinded analysis of immunofluorescence images was deposited to Zenodo via GitHub: DOI: 10.5281/zenodo.6801448. Cells containing any filamentous shapes of the Alexa Fluor 488 signal were counted as “filamentous,” ones without filamentous signal but with punctate staining were counted as “punctate,” and the remaining cells with fully cytosolic signal were counted as “cytosolic.” DAPI and β-tubulin signal was used to make sure only cells containing a single nucleus were counted, avoiding cells that have not finished dividing or are multi-nuclear. Variants with a statistically significant and largest effect on inhibitor-induced LRRK2 filament formation (<10% of LRRK2 signal-positive cells after MLi-2 treatment compared to the 34.7% of MLi-2 treated wildtype LRRK2 cells) were labelled as the “Strongest impact” group, the remaining variants with a statistically significant decrease in LRRK2 filament formation (<21%) were labelled as “Moderate impact” and the remaining variants were labelled as “Microtubule binding not significantly impacted”.

### Immunoblotting data analysis

Immunoblotting data (acquired using a LI-COR CLx Western Blot imaging system) were quantified using Image Studio Lite (Version 5.2.5, RRID:SCR_013715). Quantified data were plotted with GraphPad Prism 8 (RRID:SCR_002798). For the primary screen, data from up to 6 independent biological replicates were combined (all normalized to the wildtype LRRK2 values for each replicate). For the secondary screen, data from 2 independent biological replicates (each performed in duplicate) were combined (all normalized to the wildtype LRRK2 values for each replicate). For the immunoblotting data obtained from the primary screen, outliers of LRRK2 variant activity were determined using an arbitrary cut-off of LRRK2 expression 1.9-fold higher or lower than wildtype LRRK2 expression (designated as 1) and are presented in Figure 1C and Figure 2 as open circle data points and excluded from the variant mean. LRRK2 variants that were expressed less than ∼1.9 relative to wildtype LRRK2, or greater than ∼0.53 relative to wildtype LRRK2, were considered true representations of variant activity and are presented in Figure 1C and Figure 2 as closed circles.

### Statistical analysis

Gathered data either from immunoblotting or cell counting was analysed using GraphPad Prism 8 (RRID:SCR_002798). One- or multi-way ANOVA with Dunnett’s multiple comparisons post-hoc test was used to determine statistical significance and approximate p values for each value compared to the control mean – wildtype LRRK2.

## Data Availability

All the primary data that is presented in this study has been deposited on the Zenodo data repository ((10.5281/zenodo.6401193). All plasmids and antibodies (and associated datasheets) generated at the MRC Protein Phosphorylation and Ubiquitylation Unit at the University of Dundee can be requested through our website https://mrcppureagents.dundee.ac.uk/.

## Acknowledgements

We thank Axel Knebel for purifying recombinant Rab8A, Natalia Shpiro for generating MLi-2, and Suzanne Pfeffer for helpful discussions. We also thank the excellent technical support of the MRC protein phosphorylation and ubiquitylation unit (PPU) DNA sequencing service (coordinated by Gary Hunter), the MRC-PPU tissue culture team (coordinated by Edwin Allen), MRC-PPU Reagents and Services antibody and protein purification teams (coordinated by Dr James Hastie). This research was funded in part by Aligning Science Across Parkinson’s [ASAP-000463] through the Michael J. Fox Foundation for Parkinson’s Research (MJFF). For the purpose of open access, the authors have applied a CC BY public copyright license to all Author Accepted Manuscripts arising from this submission.

## Author contributions

A.F.K., E.P., F.T. planned and performed all experimental work, analysed data, prepared figures, and helped write manuscript. S.V.M performed the structural analysis related to Fig 7, 8 and SFig 10. M.L. performed most of the cloning required to generate LRRK2 variants. A.R.P. undertook microscopy studies for the microtubule binding analysis. S.P. and E.S. provided advice and helped with the selection of LRRK2 variants. D.R.A contributed to conceptualization of project, data analysis, writing manuscript and supervised the work.

---

**Supplementary Figure 1.**
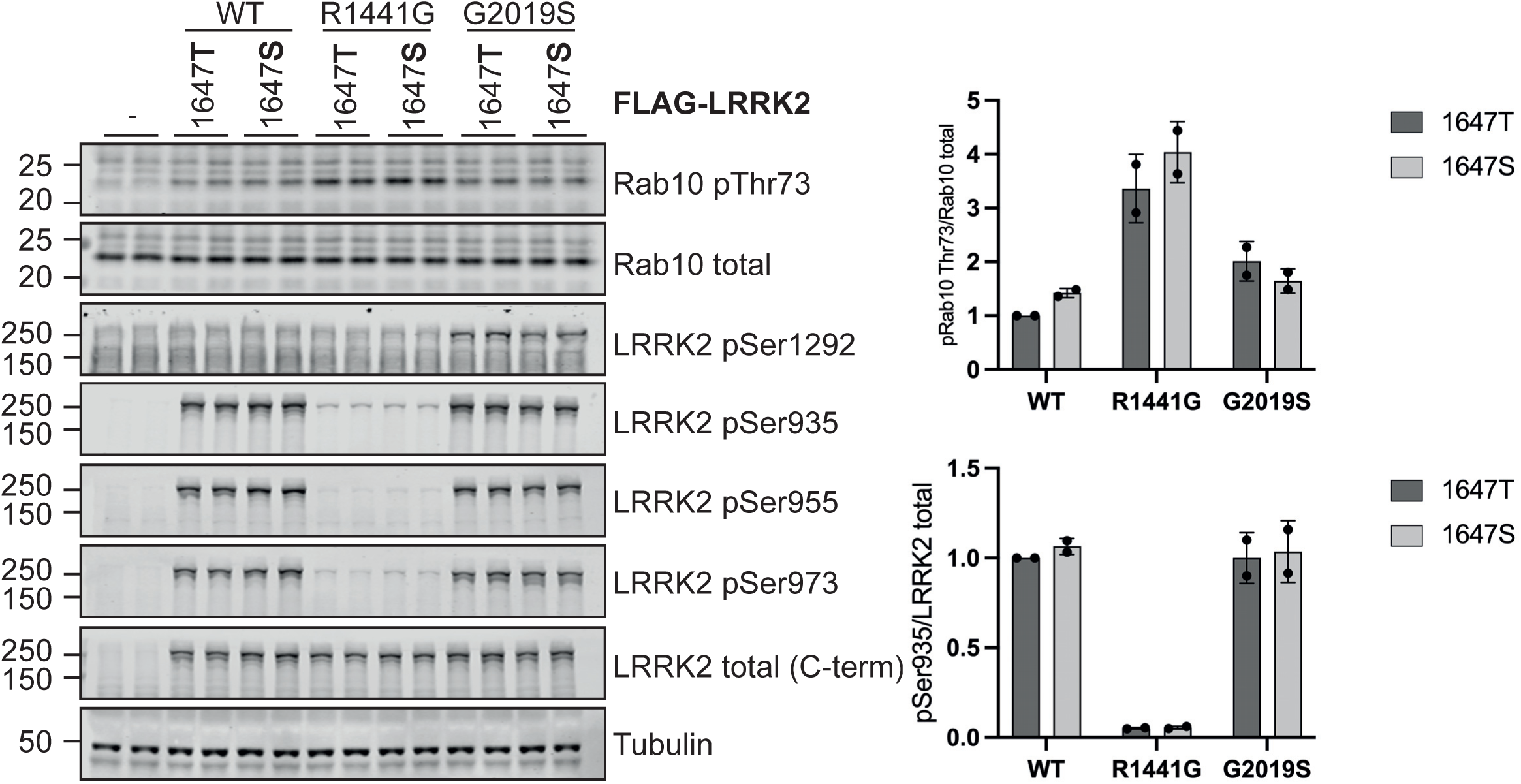
LRRK2 S1647T does not impact wildtype or pathogenic R1441G or G2019S LRRK2 activity. FLAG empty vector, FLAG-tagged LRRK2 wildtype, and the indicated variants were expressed in HEK293 cells. Cells were lysed 24 hours post-transfection and were analysed by quantitative immunoblotting with the indicated antibodies. Each lane represents a different dish of cells. The ratios of phospho-Rab10 Thr73/total Rab10 and phospho-LRRK2 Ser935/total LRRK2 were normalized to wildtype LRRK2 values. Quantified data are presented as mean ± SD and are representative of two independent experiments.

**Supplementary Figure 2.**
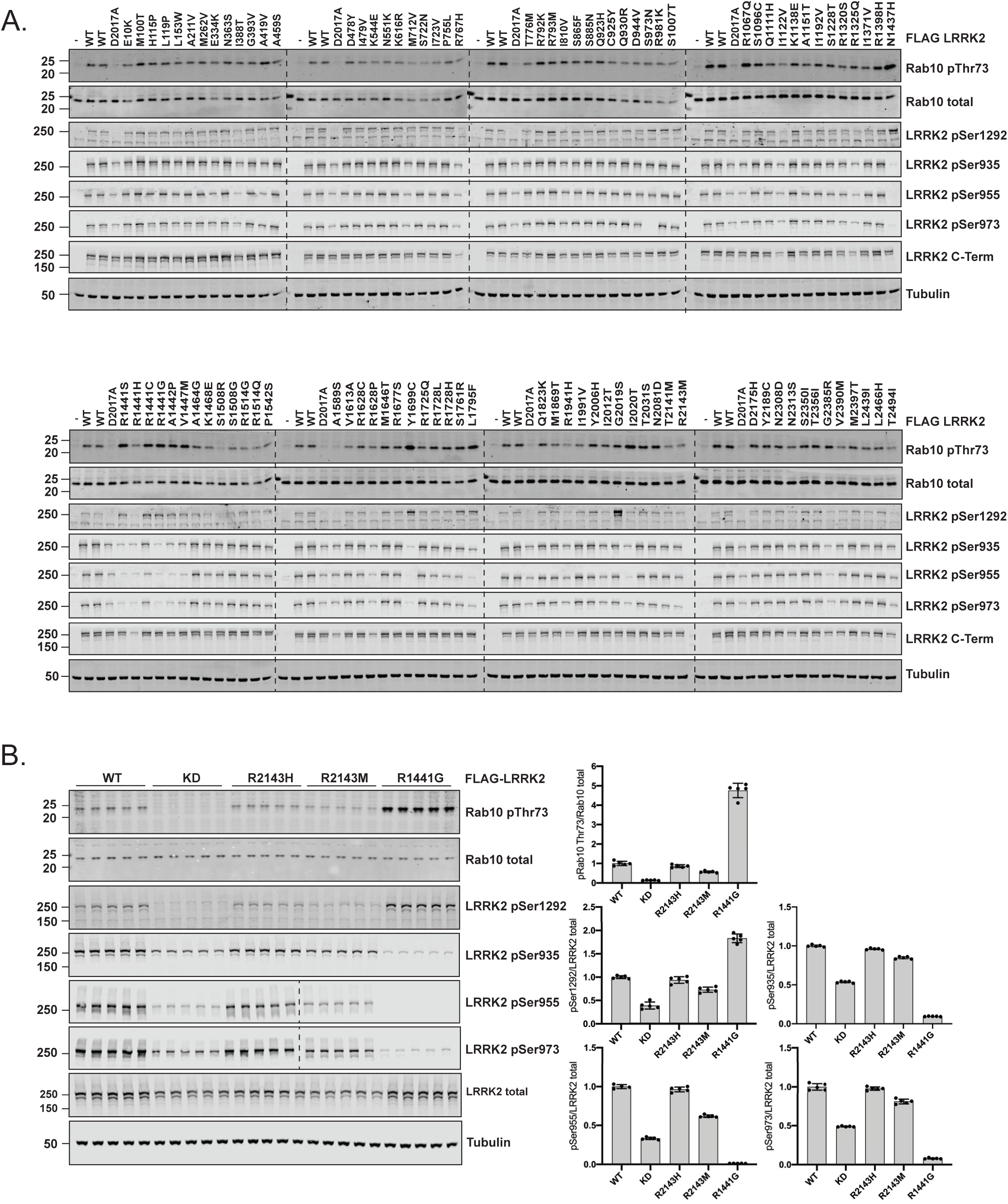
Primary quantitative immunoblot screen to assess activity of selected PD and CD-associated LRRK2 variants. (A) FLAG-tagged LRRK2 wildtype, kinase dead (KD = D2017A) and the indicated LRRK2 variants were transiently expressed in HEK293 cells. Cells were lysed 24 hours post-transfection and analysed by quantitative immunoblotting using the indicated antibodies. Immunoblot figure is representative of 6 independent biological replicates. Quantification of the combined immunoblotting data from all replicates is presented in Fig 1C, Figure 2A, Figure 2B and Figure 2C. (B) FLAG-tagged LRRK2 wildtype, kinase dead (KD = D2017A) and the indicated variants were transiently expressed in HEK293 cells. Each lane represents a different dish of cells. Cell lysate was analysed by quantitative immunoblotting with the indicated antibodies. The ratios of phospho-Rab10 Thr73/total Rab10, phospho-LRRK2 Ser935-955-973/total LRRK2, phospho-LRRK2 Ser1292/total LRRK2, were normalized to the average of wildtype LRRK2 values. Quantified data are presented as mean ± SD.

**Supplementary Figure 3.**
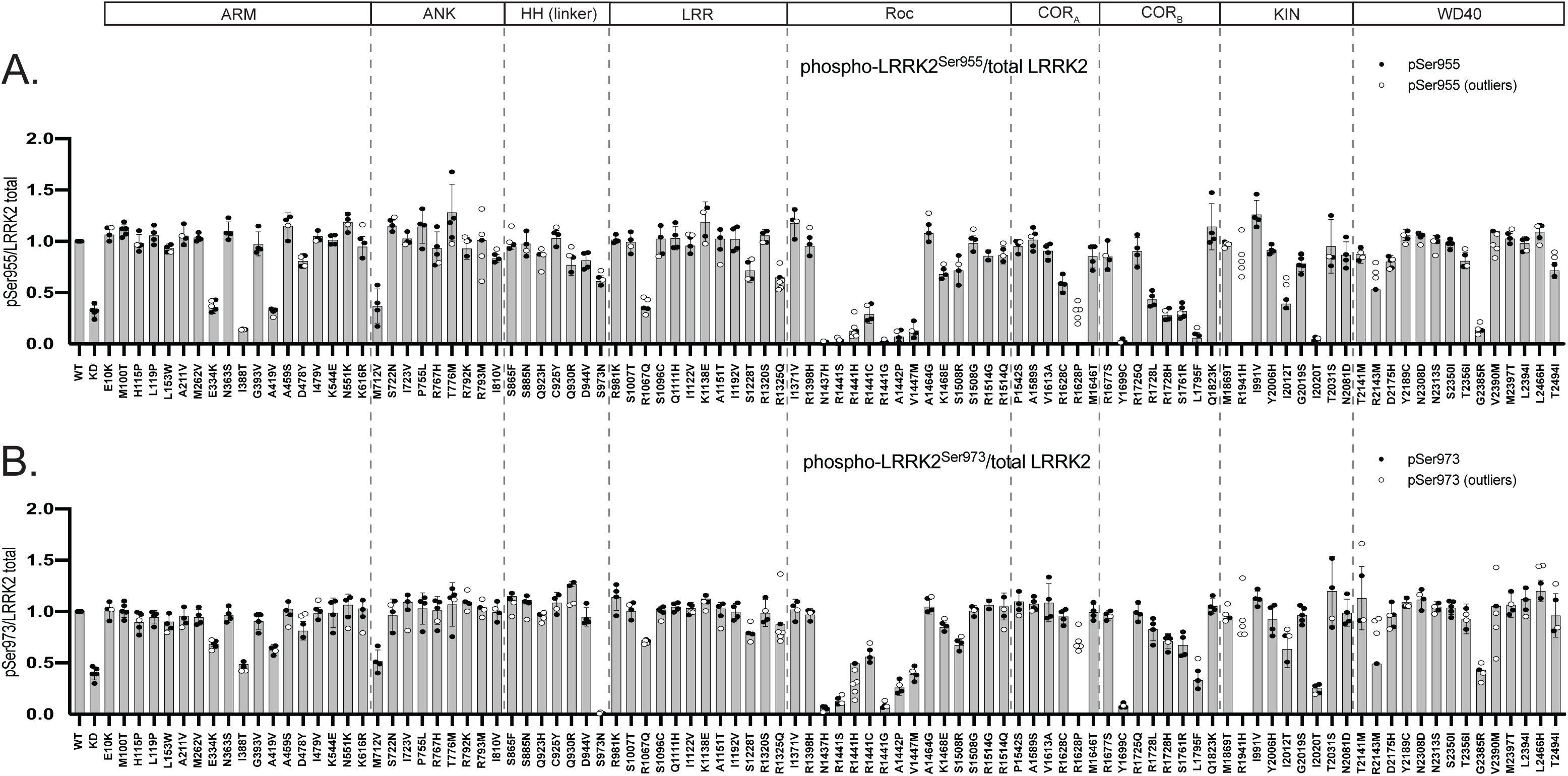
Quantitative analysis of biomarker phosphorylation of selected PD and CD-associated LRRK2 variants assessed in primary immunoblot screens. FLAG-tagged LRRK2 wildtype, kinase dead (KD = D2017A) and the indicated LRRK2 variants were transiently expressed in HEK293 cells. Cells were lysed 24 hours post-transfection and analysed by quantitative immunoblotting. Quantified immunoblotting data are presented as the ratio of LRRK2 pSer955/total LRRK2 (A) and LRRK2 pSer973/total LRRK2 (B), normalized to the average of LRRK2 wildtype values for each replicate (mean ± SD). Dashed lines segment the graphs into the corresponding regions of LRRK2 as listed in the domain schematic above panel.

**Supplementary Figure 4.**
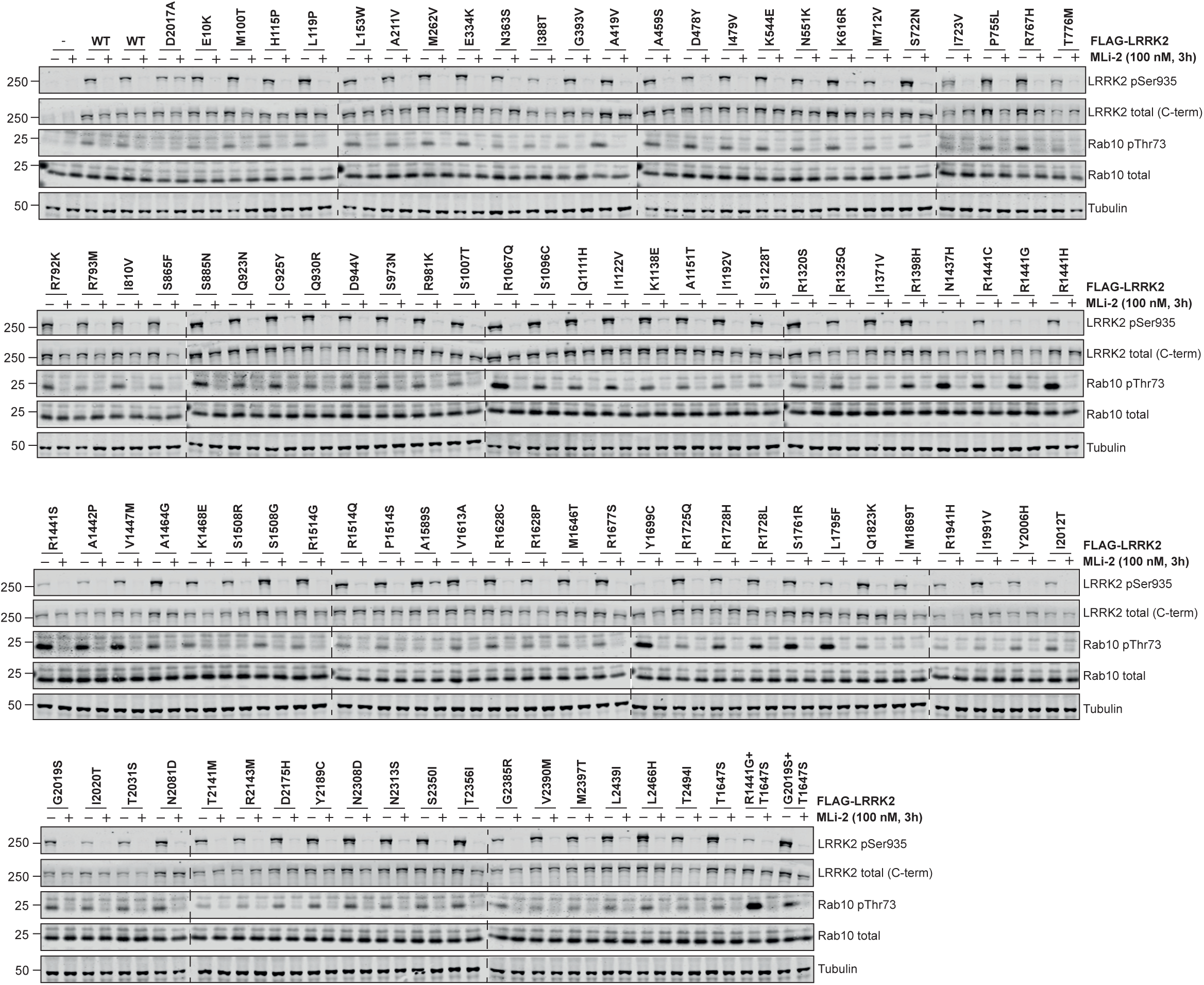
Rab10 phosphorylation mediated by selected PD and CD LRRK2 variants is reduced with MLi-2 LRRK2 inhibitor. FLAG-tagged LRRK2 wildtype, kinase dead (KD = D2017A) and the indicated LRRK2 variants were transiently expressed in HEK293 cells for 24 hours. Three hours prior to cell lysis, cells were treated with vehicle (0.1% v/v DMSO) or 100 nM MLi-2. Cell lysates were subjected to quantitative immunoblotting with the indicated antibodies.

**Supplementary Figure 5.**
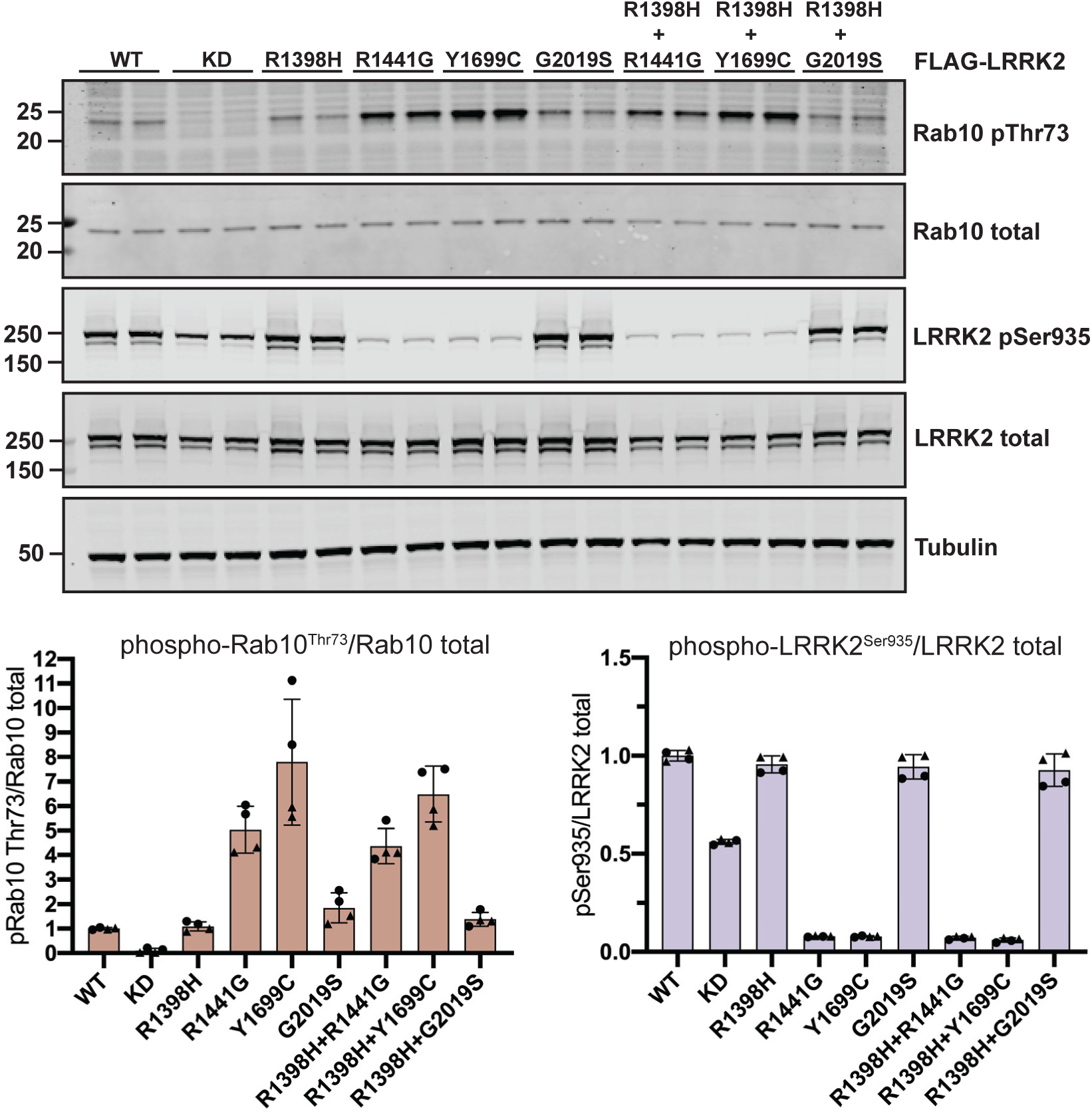
The protective LRRK2 R1398H variant does not impact wildtype or pathogenic LRRK2 R1441G, Y1699C, or G2019S activity. FLAG-tagged LRRK2 wildtype, kinase dead (KD = D2017A) and the indicated LRRK2 variants were transiently expressed in HEK293 cells for 24 hours. Each lane represents a different dish of cells. Cell lysates were subjected to quantitative immunoblotting with the indicated antibodies. Quantified immunoblotting data are presented as the ratios of phospho-Rab10 Thr73/total Rab10 and phospho-LRRK2 Ser935/total LRRK2, normalized to the average of wildtype LRRK2 values (mean ± SD). Quantified data are representative of two independent experiments.

**Supplementary Figure 6.**
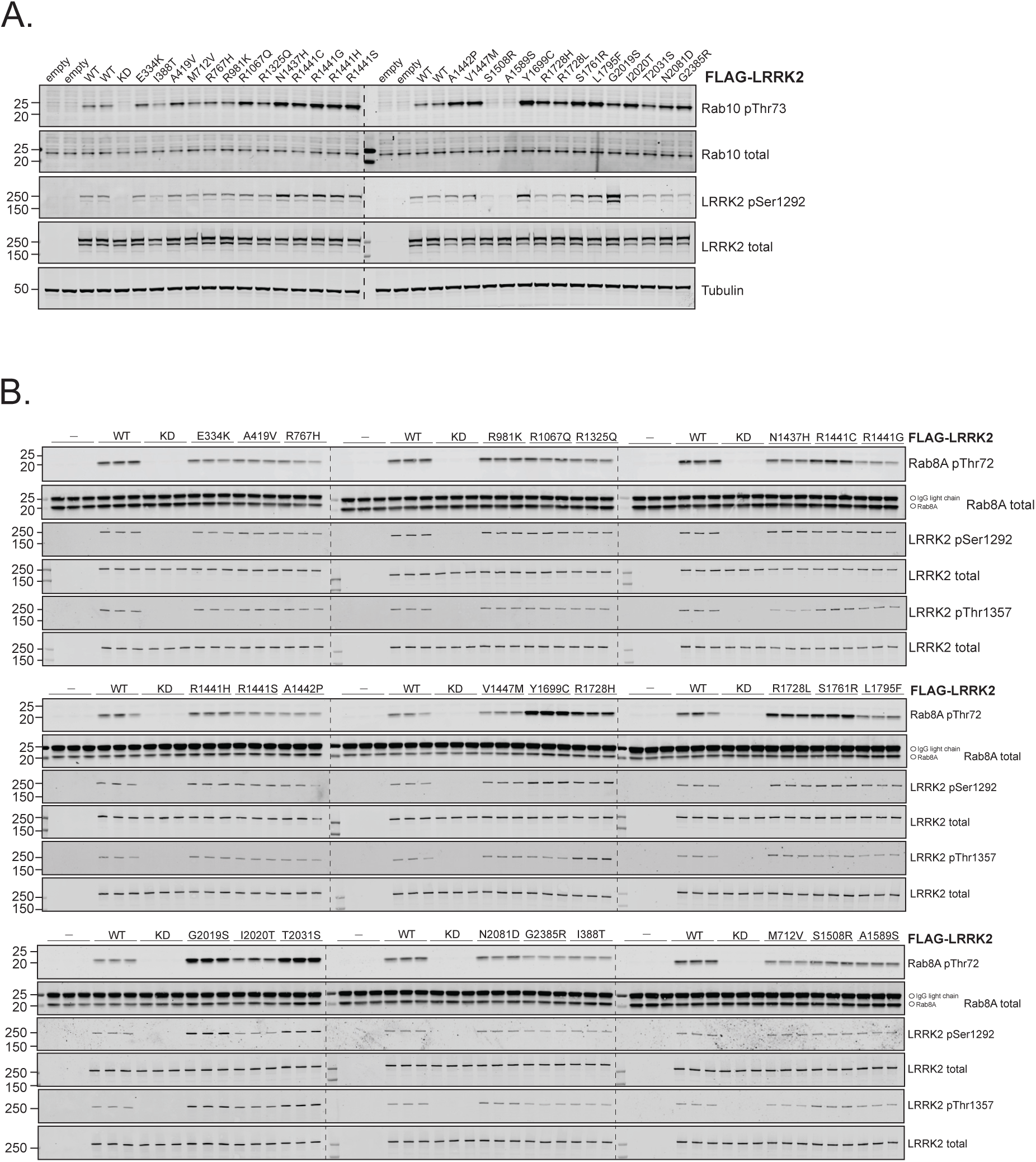
Analysis of *in vitro* LRRK2 kinase activity against recombinant Rab8A of selected LRRK2 variants. FLAG-tagged LRRK2 wildtype, kinase dead (KD = D2017A) and the indicated LRRK2 variants were transiently expressed in HEK293 cells for 24 hours. (A) Whole cell lysates were analysed by quantitative immunoblotting using the indicated antibodies. (B) FLAG-LRRK2 was immunoprecipitated from whole cell lysates and subjected to an *in vitro* kinase reaction in the presence of recombinant Rab8A. Kinase reaction products were analysed by quantitative immunoblotting using the indicated antibodies.

**Supplementary Figure 7.**
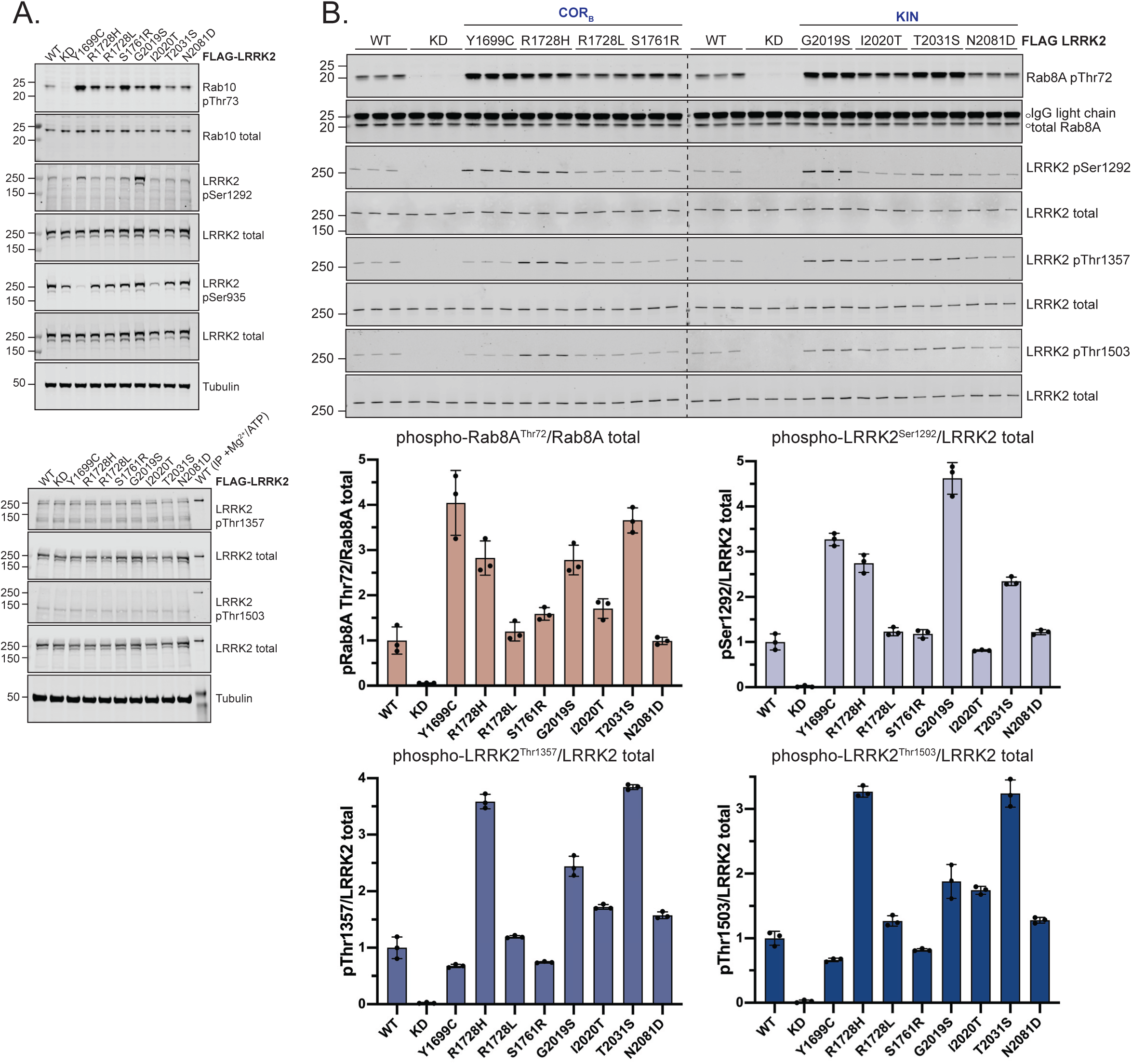
COR_B_ and kinase domain LRRK2 variants reproducibly enhance *in vitro* LRRK2 kinase activity. FLAG-tagged LRRK2 wildtype, kinase dead (KD = D2017A) and the indicated LRRK2 variants were transiently expressed in HEK293 cells for 24 hours. (A) Whole cell lysates were analysed by quantitative immunoblotting using the indicated antibodies. (B) FLAG-LRRK2 was immunoprecipitated from whole cell lysates and subjected to an *in vitro* kinase reaction in the presence of recombinant Rab8A. Kinase reaction products were analysed by quantitative immunoblotting using the indicated antibodies. Quantified immunoblotting data are presented as ratios of phospho-LRRK2 Thr1357/total LRRK2, phospho-LRRK2 Thr1503/total LRRK2, or phospho-Rab8A Thr72/total Rab8A, normalized to the average of wildtype LRRK2 values (mean ± SD).

**Supplementary Figure 8.**
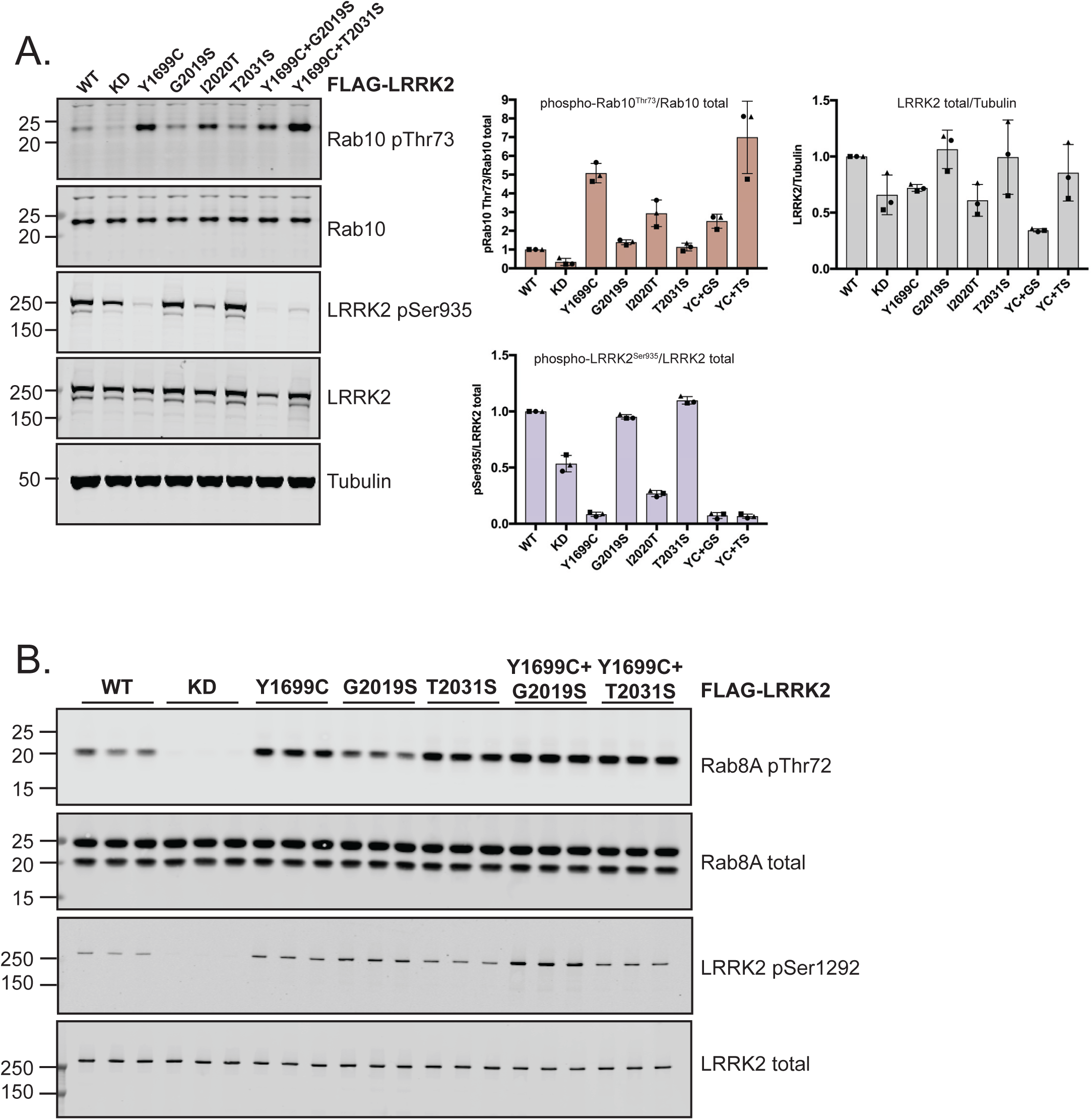
Combination of COR_B_ and kinase domain LRRK2 mutations further enhances *in vitro* LRRK2 kinase activity. FLAG-tagged LRRK2 wildtype, kinase dead (KD = D2017A) and the indicated LRRK2 variants were transiently expressed in HEK293 cells for 24 hours. (A) Whole cell lysates were analysed by quantitative immunoblotting using the indicated antibodies. Quantified immunoblotting data are presented as ratios of phospho-LRRK2 Ser935/total LRRK2, phospho-Rab10 Thr73/total Rab10, and total LRRK2/Tubulin. (B) FLAG-LRRK2 was immunoprecipitated from whole cell lysates and subjected to an *in vitro* kinase reaction in the presence of recombinant Rab8A. Kinase reaction products were analysed by quantitative immunoblotting using the indicated antibodies. Quantified data are presented in Fig 4E.

**Supplementary Figure 9.**
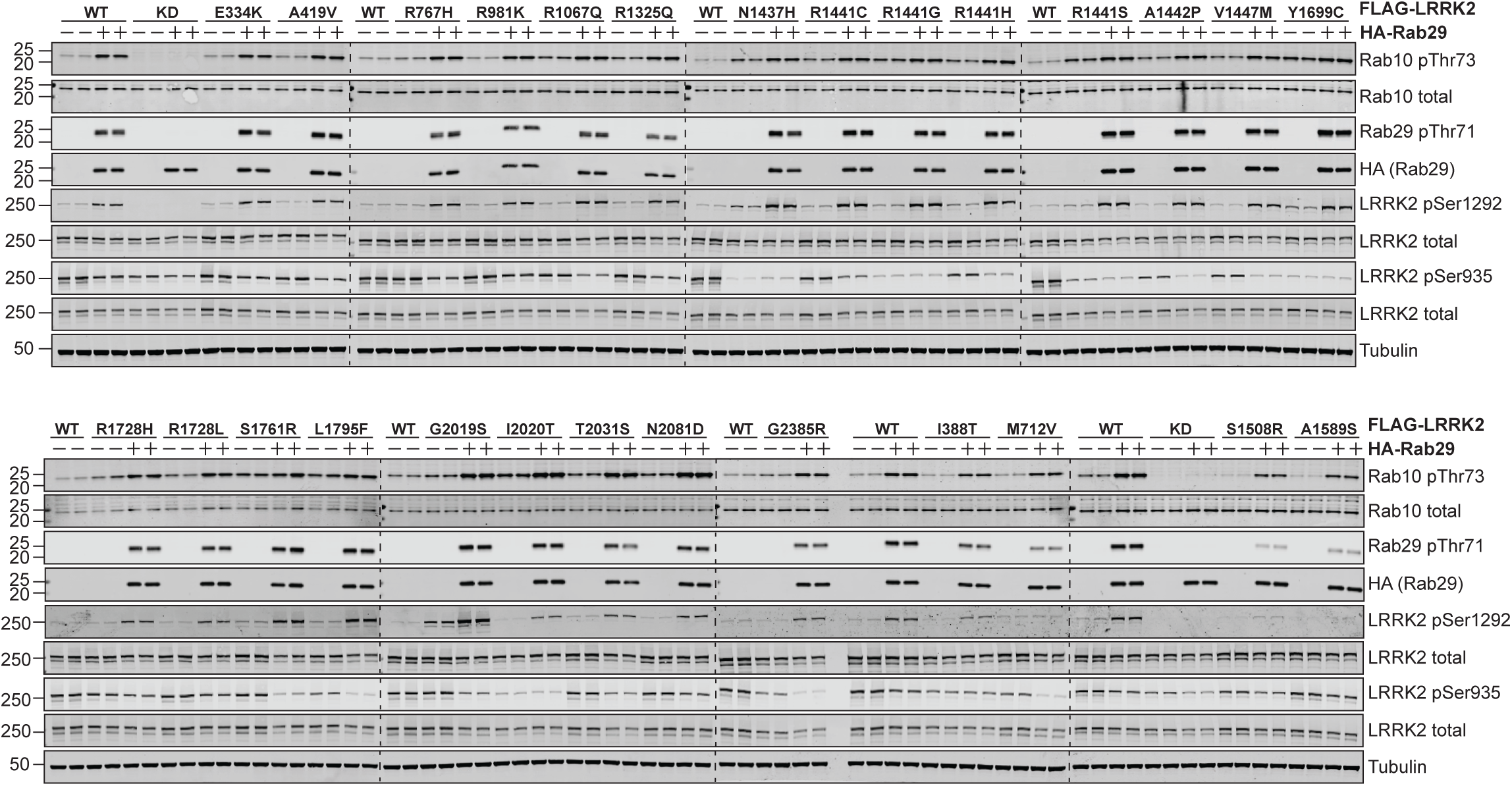
Activation of selected LRRK2 variants by Rab29. FLAG-tagged LRRK2 wildtype, kinase dead (KD = D2017A) and the indicated variants were transiently expressed in HEK293 cells with HA empty vector or HA-tagged Rab29. 24 hours post-transfection, cells were lysed and analysed by quantitative immunoblotting with the indicated antibodies. Quantified data are presented in Figure 5.

**Supplementary Figure 10.**
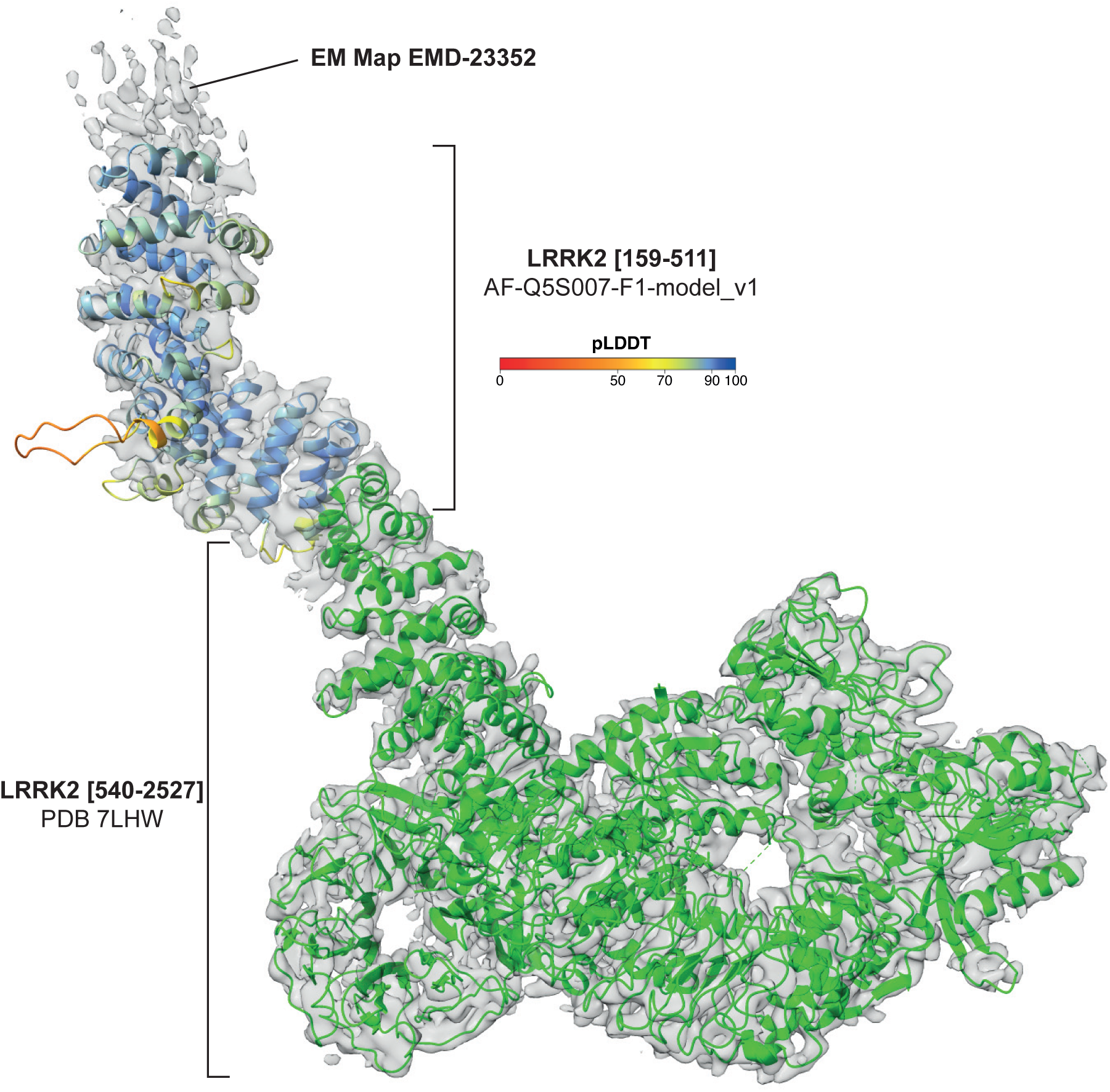
Superposition of full-length LRRK2 electron density map (PDB 7LHW) on Alphafold LRRK2 model. Alphafold model of LRRK2 ARM domain has a high local confidence score (pLDDT) and agrees well with experimental data. The ARM domain of the Alphafold model of LRRK2 (residues 159-511, AFDB AF-Q5S007-F1-model_v1) was coloured by pLDDT and fitted into the experimental cryo-EM map of full-length LRRK2 (grey, EMD-23352) using the UCSF ChimeraX “Fit in Map” tool. The remaining C-terminal LRRK2 residues [540–2527] are shown in green (PDB 7LHW

**Supplementary Table 1.**
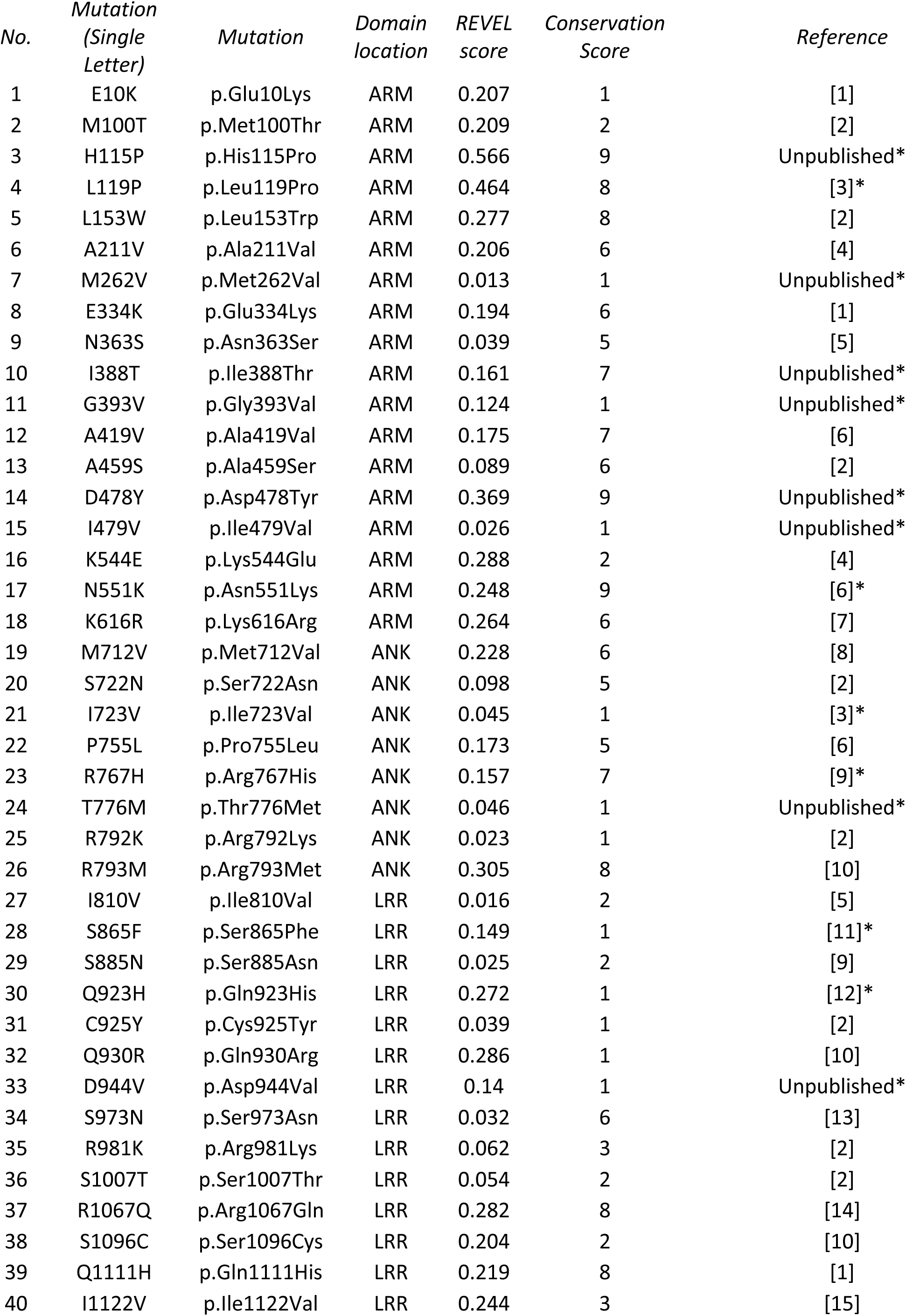
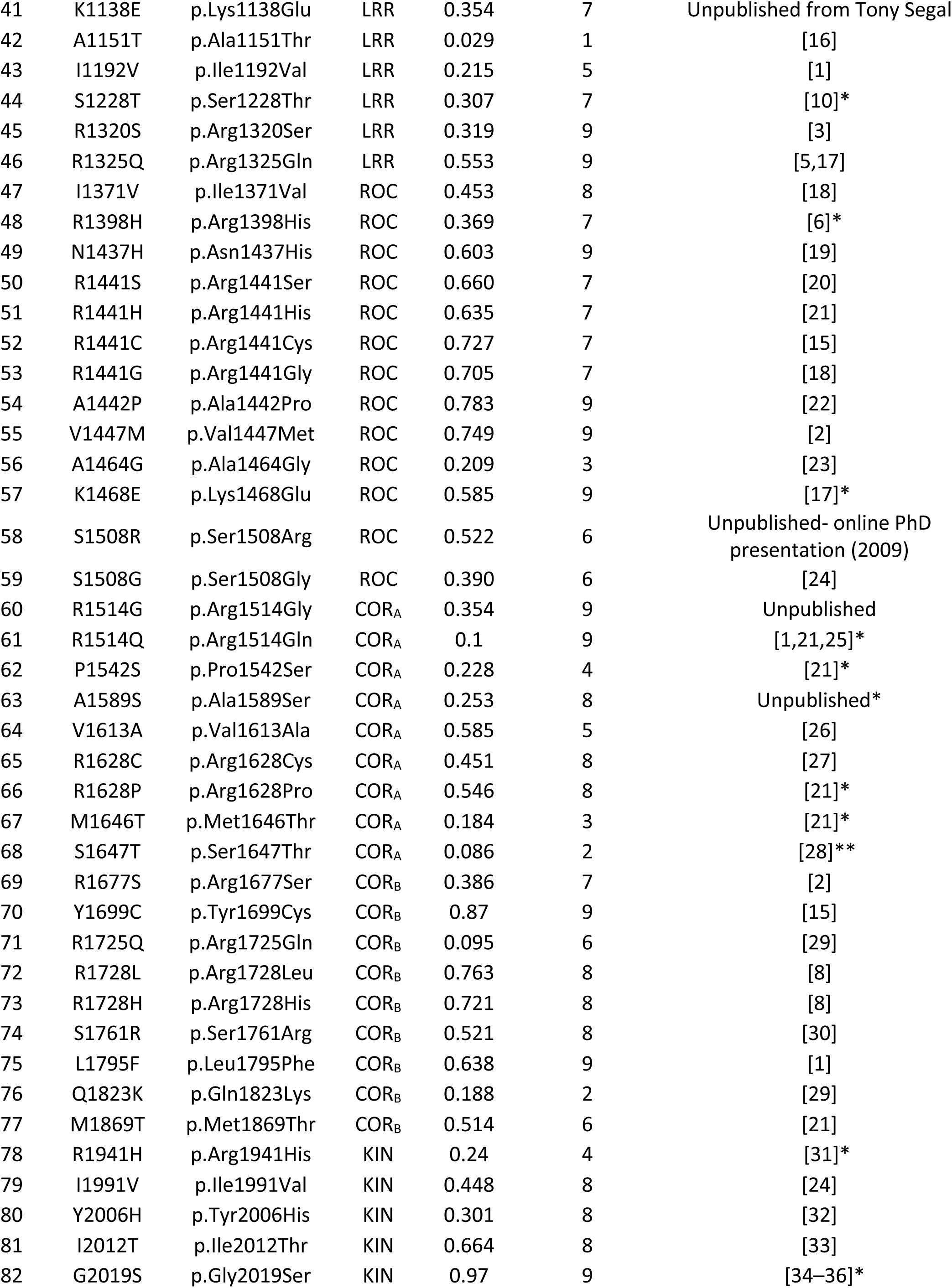
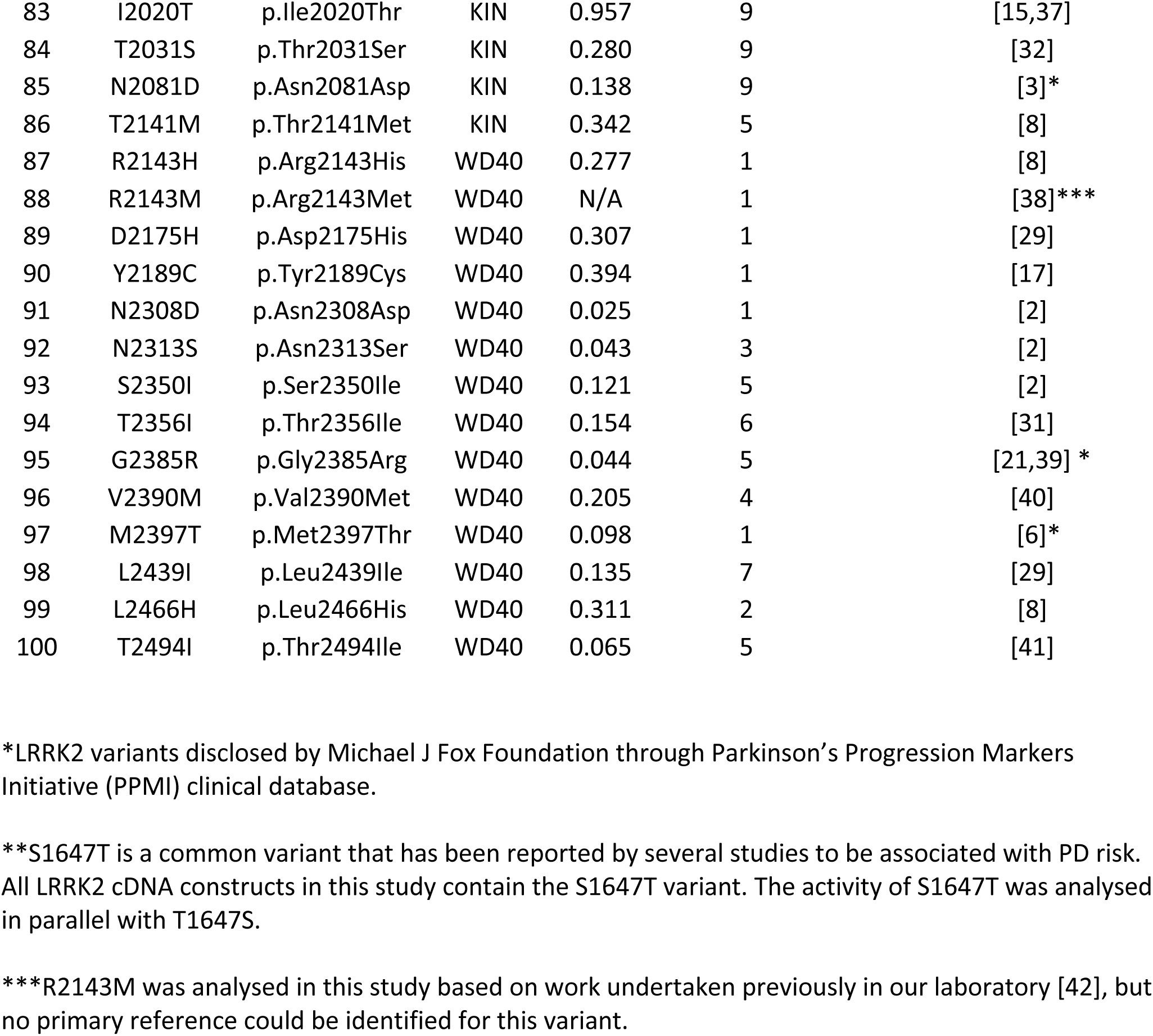
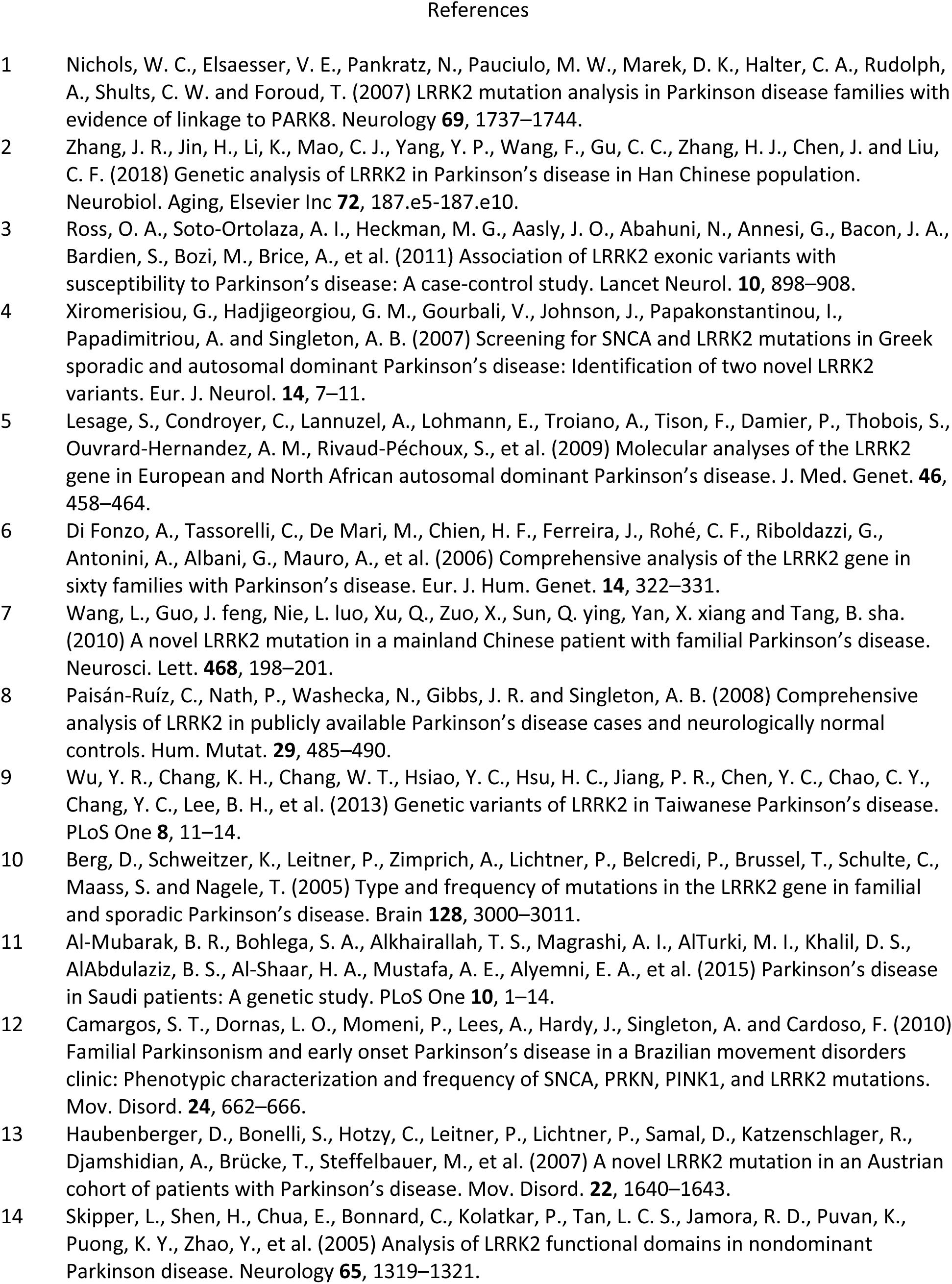
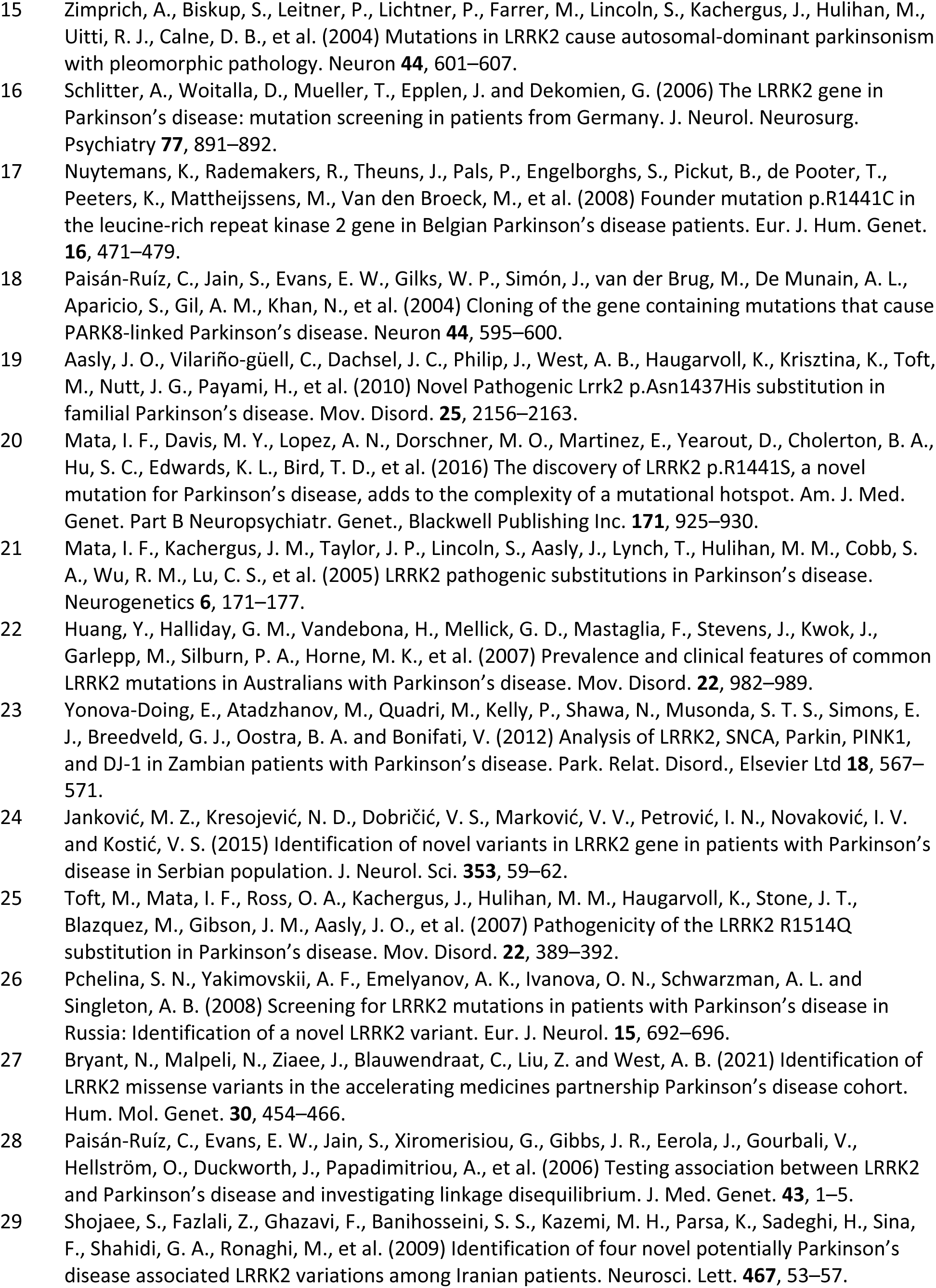
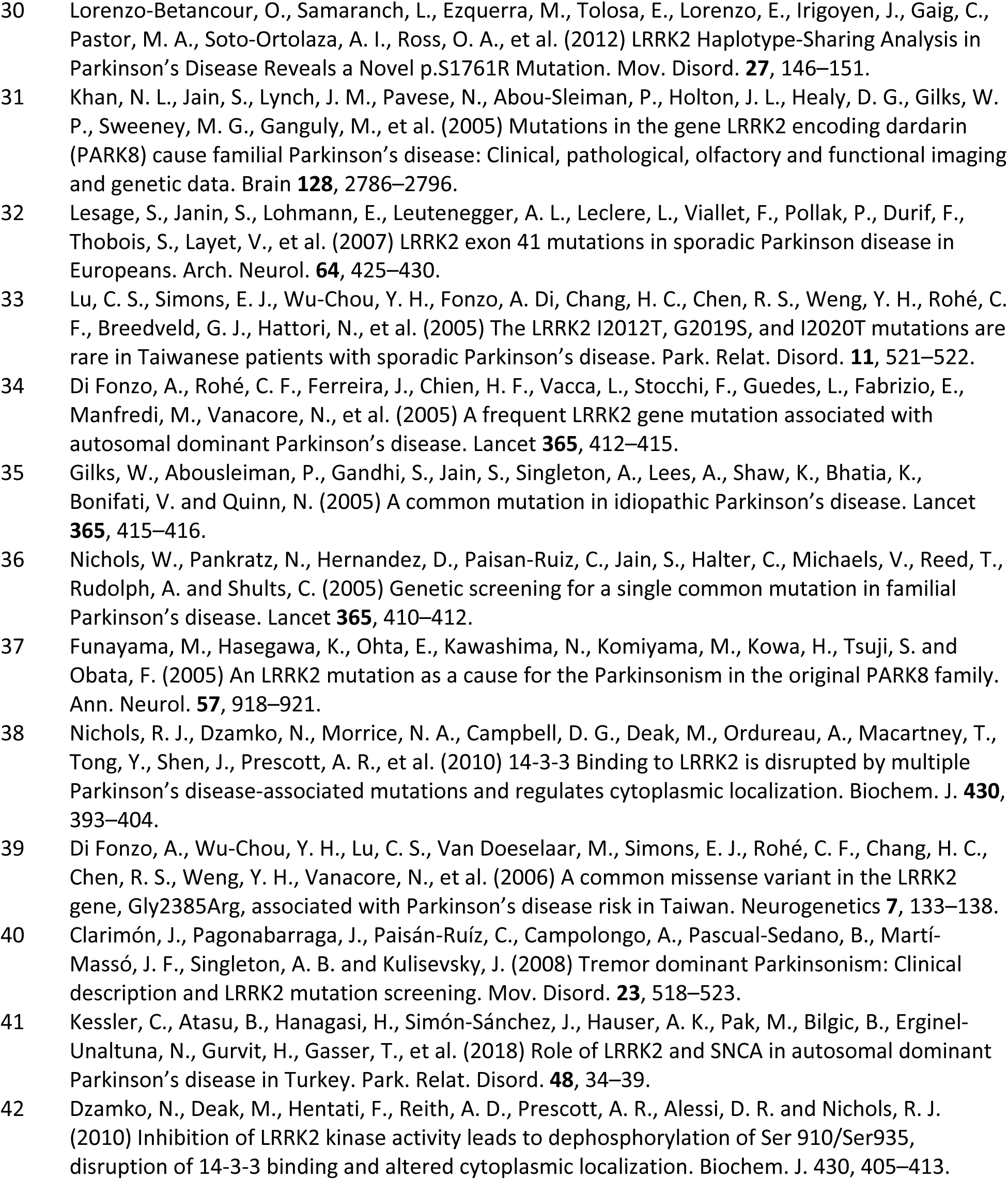

